# Nuclear lamin A/C promotes cancer cell survival and lung metastasis without restricting transendothelial migration

**DOI:** 10.1101/2020.06.23.167130

**Authors:** Francesco Roncato, Ofer Regev, Sara W. Feigelson, Sandeep Kumar Yadav, Lukasz Kaczmarczyk, Nehora Levi, Diana Drago-Garcia, Samuel Ovadia, Marina Kizner, Yoseph Addadi, João C. Sabino, Yossi Ovadya, Sérgio F. de Almeida, Ester Feldmesser, Gabi Gerlitz, Ronen Alon

## Abstract

The mechanisms by which the nuclear lamina of tumor cells controls their migration and survival are poorly understood. Lamin A and its variant lamin C are key nuclear lamina proteins that control nucleus stiffness and chromatin conformation. Downregulation of lamin A/C levels in two metastatic lines, B16F10 melanoma and E0771 breast carcinoma, facilitated cell squeezing through rigid pores, elevated nuclear deformability and reduced heterochromatin. Unexpectedly, the transendothelial migration of both cancer cells in vitro and in vivo, through lung capillaries, was not elevated by lamin A/C knockdown. Both cancer cells with lamin A/C knockdown grew normally in primary tumors and in vitro on rigid surfaces. Strikingly, however, both lamin A/C deficient melanoma and breast cancer cells grew poorly in 3D spheroids expanded in soft agar cultures. Experimental lung metastasis of both lamin A/C knockdown cells was also markedly reduced. Taken together, our results suggest that high content of lamin A/C in multiple cancer cells promotes cancer cell survival and ability to generate lung metastasis without compromising cancer cell emigration from lung vessels.

## Introduction

The nucleus is the largest and stiffest organelle in all cells and therefore imposes the main barrier for cell crossing of cellular and mechanically resistant extracellular barriers (Burke & Stewart, 2013; Friedl, Wolf, & Lammerding, 2011; Fruleux & Hawkins, 2016). The nucleus must undergo various shape changes during cell migration through cellular and extracellular barriers (Friedl et al., 2011). Lamin A and its splice variant lamin C are key nuclear lamina intermediate filament proteins that control nucleus stiffness (Burke & Stewart, 2013; Lammerding et al., 2006; Roman et al., 2017; Shin et al., 2013), and regulate chromatin conformation and accessibility (Bronshtein et al., 2015; Towbin, Meister, & Gasser, 2009). A-type lamins also control nuclear crosstalk with all types of the cell cytoskeleton, including microtubules and actin filaments (Chambliss et al., 2013; Chang, Folker, Worman, & Gundersen, 2013; Etienne-Manneville & Lammerding, 2017; Khatau et al., 2009; Lombardi & Lammerding, 2011), and thereby regulate nuclear location and response to mechanical signals from the extracellular environment (Borrego-Pinto et al., 2012; Gonzalez-Granado et al., 2014; Graham et al., 2018; Kim & Wirtz, 2015; Kirby & Lammerding, 2018; Swift et al., 2013).

Alterations in nuclear lamina stiffness can take place by various DNA damaging processes including the recently described nuclear autophagy, nucleophagy (Dou et al., 2015) or by genetic changes (J. L. V. Broers & Ramaekers, 2014). The latter changes can be experimentally introduced by controlled suppression or overexpression of lamin A/C and such changes have been studied in different types of cells migrating through variably rigid confinements in vitro and in vivo (C. M. Denais et al., 2016). The soft nuclei of most leukocytes contain low levels of lamin A/C and high levels of other lamins, primarily of the B type (Shin et al., 2013). The low ratio of A and B lamins allows leukocytes to undergo massive and rapid deformation during fast squeezing through vascular endothelial junctions and collagenous interstitial spaces (Yadav et al., 2018). How nuclear squeezing is regulated in solid cancer cells migrating through variable interstitial ECM barriers and constricted vascular spaces is only partially understood (C. M. Denais et al., 2016; Raab et al., 2016; Wolf et al., 2013). In contrast to leukocyte nuclei which express very low levels of lamins A/C, the nuclei of variably invasive metastatic tumor cells of mesenchymal origin is predicted to contain higher levels of these lamins and are therefore generally more stiff, potentially imposing higher restrictions on the ability of these cells to cross endothelial junctions and interstitial barriers (Cao et al., 2016). The cytoskeleton of endothelial cells which comprise the major barriers for nuclear squeezing is, however, fairly elastic and contractile (Barzilai et al., 2017; Heemskerk et al., 2016). In contrast, although collagenous barriers can undergo extensive remodeling by cell-generated forces and proteolysis (Infante et al., 2018) solid tumor cells with high lamin A/C content might less efficiently cope with these barriers than with endothelial barriers (Yadav et al., 2018). Support for this idea was recently provided by experiments with leukocytes overexpressing lamin A: a 10-fold increase in the ratio of lamin A to lamin B dramatically restricted leukocyte nuclear squeezing through rigid pores and dense collagenous barriers but was largely permissive for leukocyte transendothelial migration (Yadav et al., 2018).

Lamins A/C are also involved in chromatin conformation and epigenetics via their interactions with heterochromatin, transcriptionally repressed tightly folded chromatin tethered to the nuclear lamina (Becker, Nicetto, & Zaret, 2016; Dechat, Adam, Taimen, Shimi, & Goldman, 2010; Harr et al., 2015). Nevertheless, the direct contributions of lamins to cancer cell migration, growth, and malignancy have been in debate (J. L. V. Broers & Ramaekers, 2014). On one hand, lamin A/C expression is reduced in several solid cancers and cancer progression was suggested to correlated with lower lamin A/C expression (Bell & Lammerding, 2016; C. Denais & Lammerding, 2014; Kaufmann, Mabry, Jasti, & Shaper, 1991), it is absent in around 40% of human breast cancer tissues (Capo-chichi et al., 2011), and decreased lamin A/C expression is a sign of poor prognosis in skin, breast, lung and colon cancers (J. L. Broers et al., 1993; Capo-chichi et al., 2011; Venables et al., 2001). Since lamin A/C knockdown nuclei can be more easily deformed and squeeze through rigid confinements, it has been postulated that tumor cells with low expression levels of these lamins can more readily invade tissues (Davidson, Denais, Bakshi, & Lammerding, 2014). Nevertheless, the direct in vivo evidence for this assumption has never been provided. Furthermore, overexpression of A-type lamins in some cancer cells was shown to correlate with enhanced growth and faster migration via the activation of the PI3K/AKT/PTEN pathway (Kong et al., 2012). Part of these discrepancies are attributed to the complex roles of A-type lamins in protecting the nuclei from mechanical nuclear rupture, DNA damage, and cell growth arrest (Cho et al., 2019). These discrepancies have motivated us to address these standing questions by controlled downregulation of lamin A/C expression introduced into multiple bona fide metastatic cells. We specifically wished to address whether extravasation of circulating cancer cells from different blood vessels, and in particular from the relatively impermeable lung capillaries, are favored by the softening of tumor cell nuclei via downregulation of lamin A/C. We hence systematically addressed both in vitro and in vivo in syngeneic mice models if the forced downregulation of lamins A/C expression with retained levels of B lamins alters both invasive and proliferative properties of two prototypic metastatic cell lines, namely, B16 melanoma and E0771 breast carcinoma. In vitro, as expected, downregulated lamin A/C levels introduced by ectopic expression of lamin A/C specific shRNA dramatically facilitated the squeezing of both cells through rigid pores, reduced their heterochromatin content, and altered their transcriptional signature. Surprisingly, however, circulating lamin A/C knockdown cells normally emigrated from the pulmonary circulation and normally accumulated inside the lung parenchyma. Nevertheless, the ability of lamin A/C low melanoma and breast cancer cells to generate metastatic lesions in the lungs was markedly compromised. In vitro, both B16 melanoma and E0771 breast carcinoma cells with lamin A/C knockdown exhibited specialized proliferation defects when grown in spheroids within 3D environment whereas their intrinsic growth on solid 2D surfaces remained intact. Our results collectively suggest that high nuclear lamin A/C content does not compromise the squeezing ability of melanoma and breast cancer cells through physiological endothelial barriers. High content of lamin A/C is necessary, however, for optimal growth of these multiple cancer cells in multi-cellular assemblies in vitro and for their ability to generate lung metastasis in vivo.

## Results

### Lamin A downregulation in B16F10 melanoma increases nucleus deformability and squeezing through rigid pores but does not affect transendothelial migration

To gain insight into the contribution of type A lamins to the ability of solid tumor cells to translocate their nuclei across endothelial barriers, we downregulated lamin A and lamin C expression in the nuclei of the B16F10 melanoma cells by stably introducing a lamin A/C shRNA construct targeting exon 8 of the *Lmna* gene (Figure 1A, B). Since the nuclear lamina stiffness is sensitive to the ratio between type A and type B lamins in both mesenchymal and hematopoietic cells (Shin et al., 2013), we expected that lamin A/C downregulation would also increase nuclear deformability and squeezing capacities in our melanoma cell model. Downregulating lamin A and lamin C expression in the B16 melanoma line by 90%, but leaving lamin B1 levels intact, altered the ratio of type A to type B lamins by approximately 10-fold (Figure 1A). As expected, lamin A/C downregulation resulted in a dramatic enhancement of B16 squeezing through small rigid pores but much less so through large pores (Figure 1C, D). A similar gain of squeezing of lamin A/C downregulated B16F10 cells was observed with another lamin A/C shRNA construct (Figure 1-figure supplement 1A) and in cells subjected to transient *Lmna* gene exon-4 targeted siRNA mediated downregulation of lamin A/C transcription (Figure 1-figure supplement 1B).

**Figure 1.**
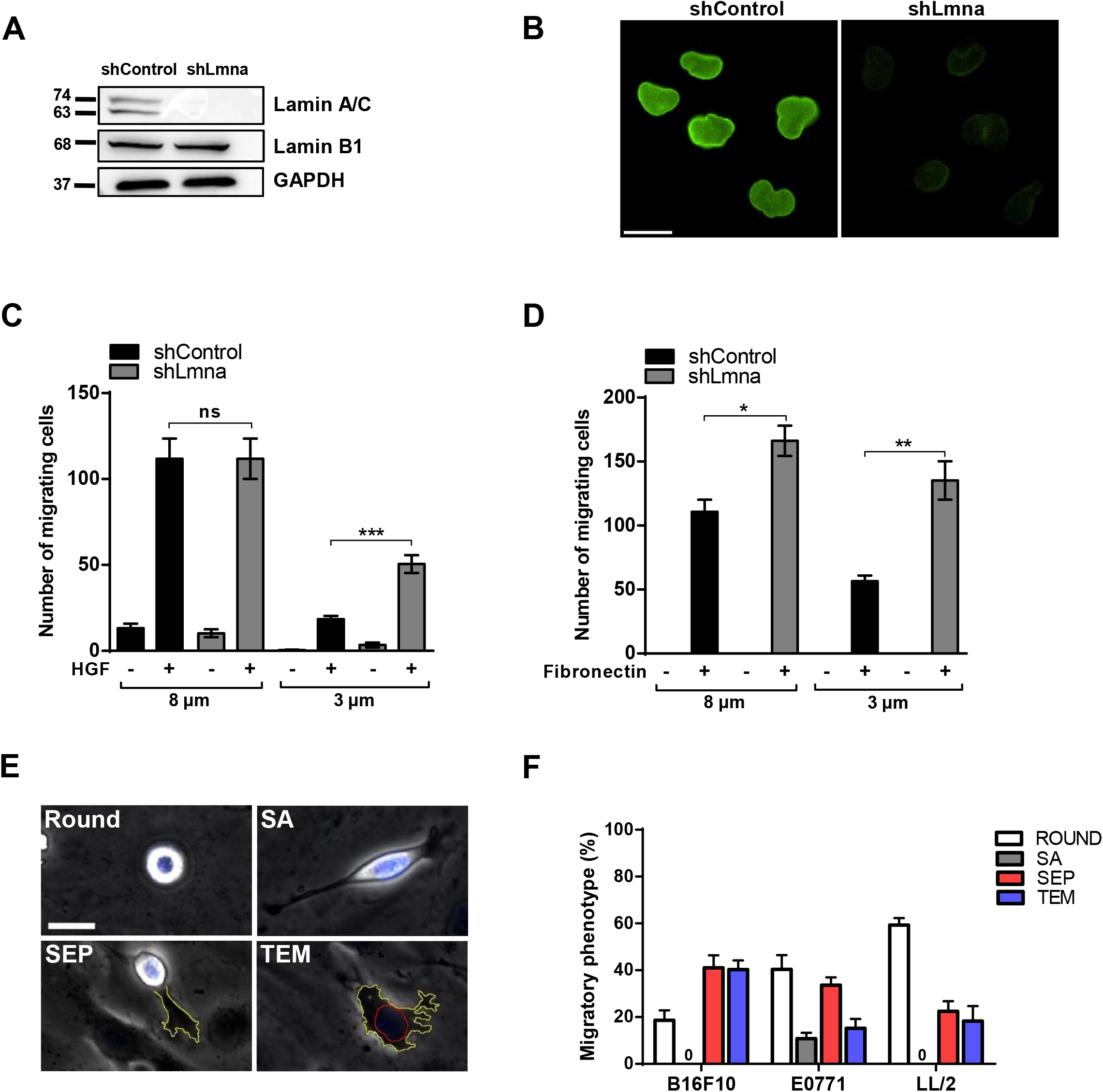
Downregulation of lamin A/C increases melanoma squeezing through small rigid pores but does not facilitate tumor transendothelial migration in vitro. (A) Expression levels of lamin A/C and lamin B1 in B16F10 cells transduced with control or lamin A/C shRNA. Glyceraldehyde-3-phosphate dehydrogenase (GAPDH) was used as loading control. (B) Immunostaining of lamin A/C (green) in B16F10 shControl and shLmna cells. Scale bar, 20 μm. (C) Chemotactic migration of B16F10 shControl and shLmna cells through 8 or 3 μm pores transwell filters in presence (+) or absence (−) of HGF (50 ng/ml) for 4 hours. Values represent the mean ± SEM of five fields in each experimental group. Results shown are from a representative experiment of three. ***p = 0.0002 (D) Haptotactic migration of B16F10 shControl and shLmna cells through 8 or 3 μm pores transwell filters in the presence or absence of fibronectin for 4 hours. Values represent the mean ± SEM of five fields in each experimental group. Results shown are from a representative experiment of three. *p = 0.0107, **p = 0.0024. (E) Representative images of distinct tumor cell categories (referred to as migratory phenotypes) taken from time-lapse videomicroscopy segments of individual B16 cells: round, spread above (SA), forming sub endothelial pseudopodia (SEP), and completing transendothelial migration (TEM). Scale bar represents 20 μm. See also Video 2. (F) Migratory phenotypes of B16F10 melanoma, E0771 breast carcinoma and LL/2 Lewis Lung Carcinoma interacting with bEnd.3 monolayers. Values represent the mean ± SEM of three fields in each experimental group (n= 40 cells). The experiment shown is representative of three.

To assess if and how these changes in the composition and deformability of the tumor nucleus promotes tumor cell squeezing through physiological endothelial barriers, we established a new in vitro video-microscopy based assay in which nuclear squeezing of tumor cell crossing confluent monolayers of bEnd.3 murine endothelial cells can be compared. Nucleus location, deformation and squeezing in individual transmigrating tumor cells could be readily tracked in real time by fluorescence and phase contrast microscopy of tumor cells whose nuclei were prelabeled with the nuclear dye Hoechst (Video 1-2). This assay allowed us to follow how individual tumor cells complete an entire sequence of TEM steps immediately after settling on confluent endothelial monolayers, including protrusion, generation of large sub-endothelial pseudopodia (lamellipodia), squeezing their nuclei, tail detachment, and locomotion underneath the endothelial monolayer (Figure 1E, Video 2). Interestingly, out of three murine cancer cell lines compared in this new assay, namely B16F10, E0771, and LL/2, B16 melanoma exhibited the highest extent of TEM (Figure 1F). Notably, the TEM of B16F10 as well as of other cancer cells compared in this system was over 30-fold slower than that of leukocytes (Barzilai et al., 2017; Yadav et al., 2018) due to a very slow formation of the sub-endothelial leading edge (t= 26±22 min for B16F10 as compared to 30±15 s for T cells), and the inability of the tumor nucleus to translocate into this leading edge as during leukocyte TEM (Barzilai et al., 2017; Yadav et al., 2018).

Strikingly, and in contrast to the squeezing results across rigid pores (Figure 1C, D), the extent of B16F10 TEM was not increased by lamin A/C knockdown (Figure 2A, B, Video 3). However, the extent of nuclear deformation for lamin A/C downregulated nuclei was significantly greater than that of the control tumor cells (Figure 2C). The nuclei of the lamin A/C knockdown B16F10 deformed more readily also when these cells spread on a non-confined 2D substrate coated with the basement membrane deposited by the endothelial monolayer, whereas the motility of lamin A/C knockdown cells remained normal (Figure 2D, Video 4). These results collectively suggest that the reduced nuclear stiffness and the increased nuclear deformability of lamin A/C knockdown B16F10 cells (Figure 2C, D) do not provide a migratory advantage for the transmigration of these cells through endothelial barriers. These observations also suggest that differences in tumor cell squeezing through infinitely rigid barriers do not correlate with squeezing through physiological cellular barriers. This finding could be due to the high mechanical flexibility of the endothelial cytoskeletal barriers maintained by their rapid actin turnover (Ofer, Mogilner, & Keren, 2011) and dynamic contractility (Heemskerk et al., 2016).

**Figure 2.**
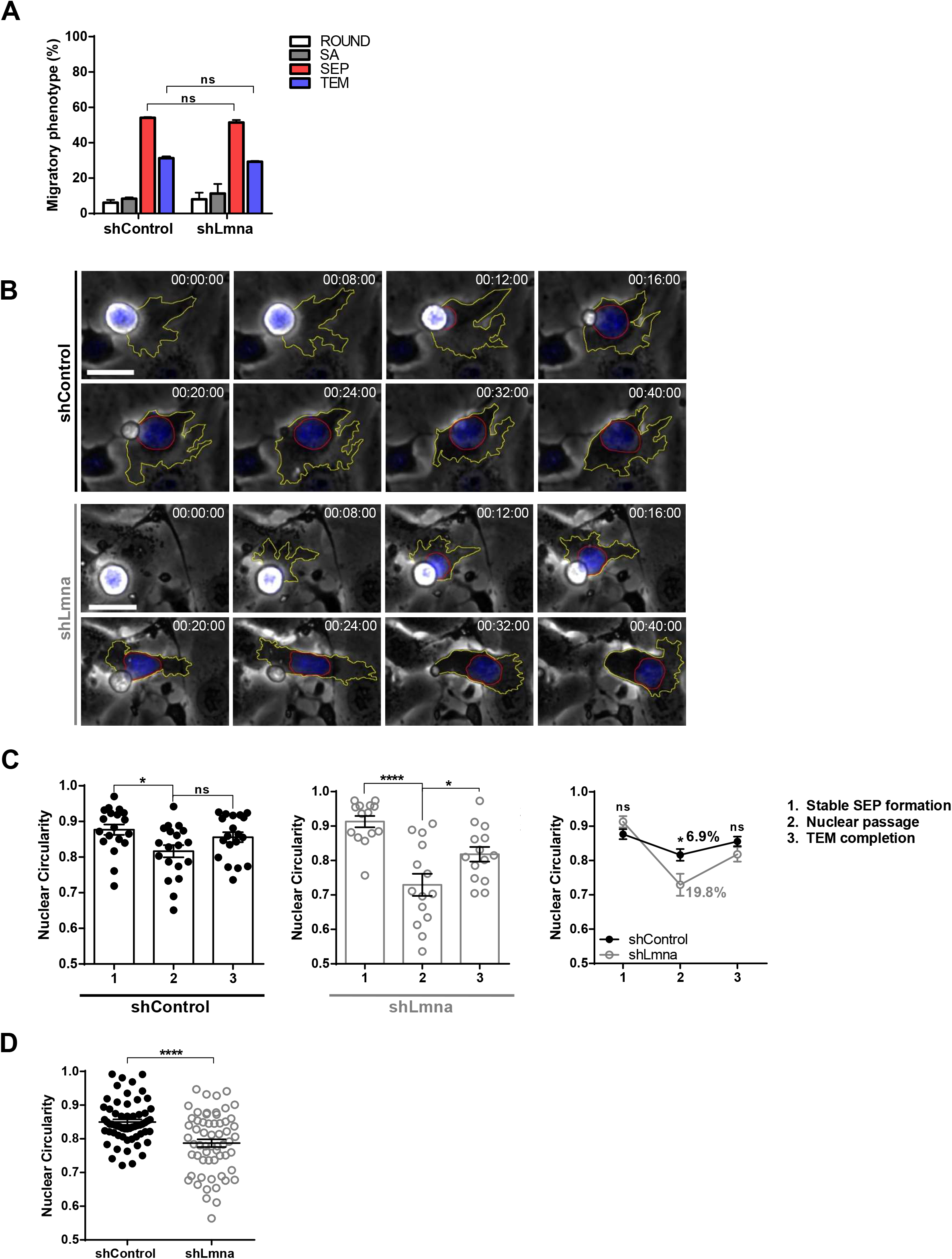
Downregulation of lamin A/C does not facilitate melanoma TEM but increases tumor nucleus deformability. (A) Migratory phenotypes of control and lamin A/C knockdown B16F10 melanoma (shControl and shLmna, respectively) interacting with unstimulated bEnd.3 cells. Values represent the mean ± SEM of four fields in each experimental group (n = 50). The experiments shown are representative of three. (B) Serial images of representative control and lamin A/C knockdown B16F10 melanoma cells, labeled with Hoechst. Time intervals are depicted in each image. The contours of the melanoma cell’s leading edge and nucleus are outlined in each image in yellow and red respectively. See also Video 3. (C) Nuclear circularity variation during the distinct indicated phases (1-3) of transendothelial migration of B16F10 shControl and shLmna cells. Values represent the mean ± SEM of six fields in each experimental group (n = 15). The experiment shown is representative of three. *p = 0.0105 (shControl); ****p < 0.0001, *p = 0.0288 (shLmna). The right panel depicts the mean nuclear circularity values for each group of B16F10 cells. The percent changes in mean circularity values are shown on top of the line plots. *p = 0.0152. (D) Nuclear circularity of B16F10 shControl and shLmna cells spread on a bEnd.3-derived basement membrane. n = 50 cells per group. Values are mean ± SEM. ****p < 0.0001. See also Video 4.

### Lamin A downregulation does not increase B16F10 extravasation across lung vessels in vivo and does not accelerate melanoma apoptosis in the lung parenchyma

Our in vitro results thus indicated that lamin A/C downregulation in B16F10 cells enhance their squeezing through rigid confinement without affecting ability to transmigrate across endothelial barriers. To assess the distinct migratory outcomes of lamin A/C downregulation in B16 melanoma cells in vivo, we introduced an experimental lung metastasis model based on i.v. injection of minute numbers of fluorescently labeled cells (Figure 2-figure supplement 1), in order to minimize non-physiological inflammatory responses associated with a bolus of cancer cells that simultaneously enter the lung vasculature. Tumor metastasis in lung spreads nearly exclusively through the pulmonary capillaries (Miles, Pruitt, van Golen, & Cooper, 2008), an extensive network of relatively impermeable capillaries (Chambers, Groom, & MacDonald, 2002) considered to be poorly permeable compared with vessels targeted by metastatic cells at other organs like the bone marrow and liver (Valastyan & Weinberg, 2011). I.V. injections of tumor cells are extensively used for studying hematogenous dissemination and expansion in the lung (Gorelik & Flavell, 2001). Since the tumor cells are introduced i.v. in a single event, their arrival at the lung vasculature is synchronized which allows accurate temporal dissection of the earliest extravasation steps taken by individual circulating metastatic cells entering the lung vasculature several hours post injection. To determine the effects of lamin A/C downregulation on earliest tumor cell extravasation across the lung capillaries, we compared the cancer cell partition inside and outside lung vessels with newly developed 3D imaging of the injected fluorescently labeled cancer cells in relation to CD31 stained lung vessels (Figure 3A, B, Video 5). Strikingly, lamin A/C downregulated B16F10 cells extravasated at similar efficiencies to normal B16F10 cells (Figure 3C, Videos 6-8). Lamin A/C downregulation also did not affect the total number of B16F10 cells accumulated inside the recipient lung at this early time points (Figure 3D). These results suggest that unlike its dramatic effects on tumor squeezing in vitro, but consistent with our TEM results in vitro (Figure 2A), lamin A/C suppression does not facilitate melanoma extravasation across lung vessels in vivo.

**Figure 3.**
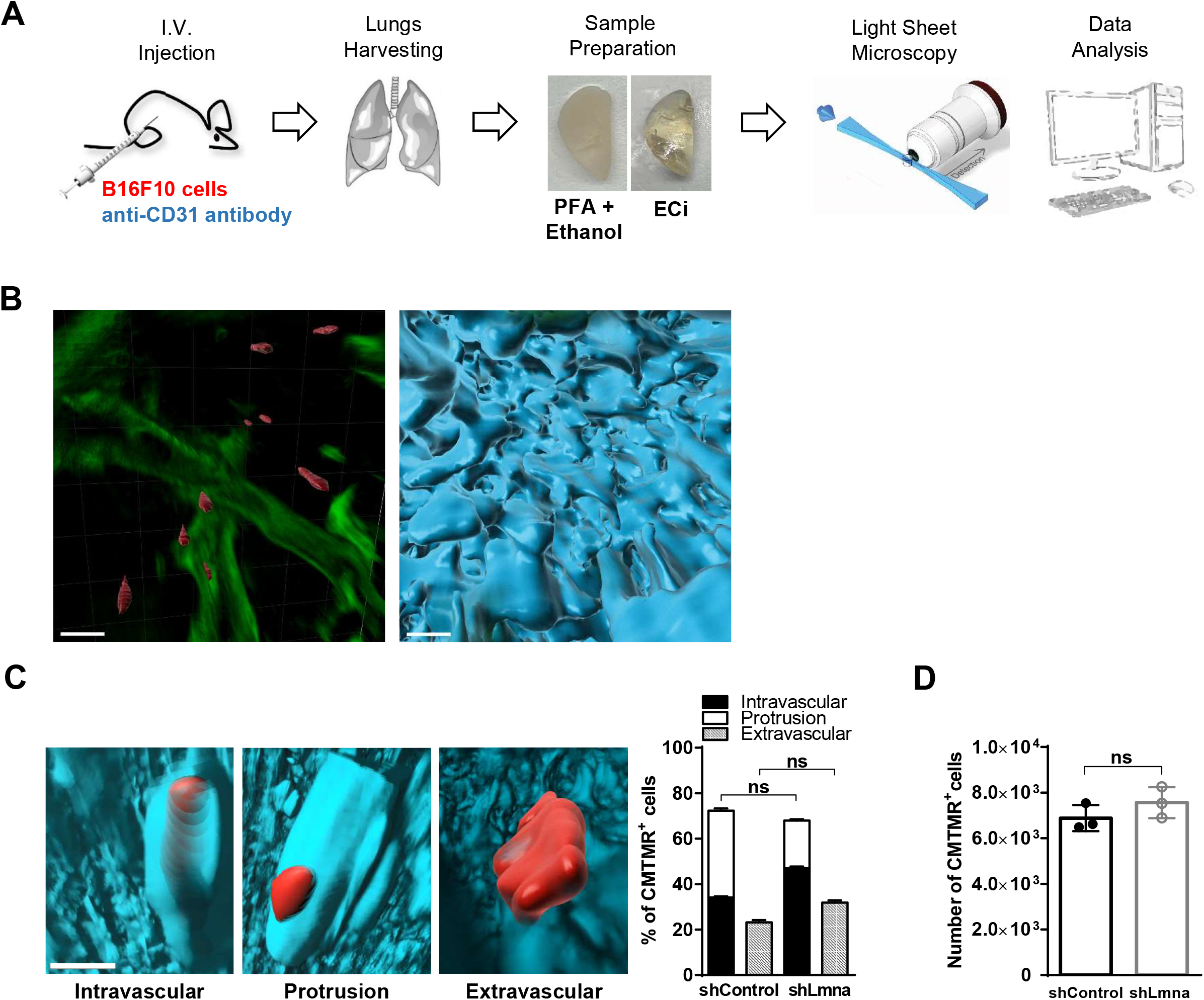
In vivo melanoma crossing of lung capillaries is not enhanced by lamin A/C downregulation. (A) Schematic representation of the LSFM analysis. (B) Visualization of 3D bronchial structures (green, autofluorescence) together with B16 melanoma cells (red, CMTMR) and alveolar capillaries (cyan, CD31). Scale bars, 100 μm. See also Video 5. (C) Representative 3D images of intravascular, extravascular, and protrusive tumor cells across the CD31-labeled lung vasculature together with the percentage of B16F10 shControl and shLmna cells present in a volume of 5×10^9^ μm^3^ of the left lung lobe (3 h after injection) counted using Imaris software. The results shown are representative of 3 experiments. Scale bar, 100 μm. See also Videos 6-8. (D) Number of CMTMR-labeled B16F10 shControl and shLmna present in the lungs of recipient mice, 3 h after i.v. injection. Data are mean ± SEM. The experiment shown is representative of two.

### Lamin A/C downregulation reduces the content of H3K9Me3 heterochromatin but does not alter DNA stability

Lmna knockout (KO) was shown to reduce the major marker of constitutive heterochromatin H3K9me3 and the methyltransferases that generate it in primary cells (Liu et al., 2013). As predicted, lamin A/C downregulation led to lower levels of H3K9Me3 and significantly reduced levels of SUV39H2 and SETDB1, two main histone methyltransferases that maintain this suppressive epigenetic marker (Figure 4A, B). In contrast, facultative heterochromatin levels probed by the H3K27me3 marker remained unchanged (Figure 4B). This global H3K9me3 reduction was not associated with global repression of transcription as determined by a transcription run-on experiment (Figure 4C). Notably, transcription of specific genes may be altered by reduced H3K9me3 levels and lamin A/C deficiency was reported to increase chromatin dynamics, which could also affect transcription (Bronshtein et al., 2015; Solovei et al., 2013; Sullivan et al., 1999). To evaluate these possibilities, we performed RNA-seq analysis on control and lamin A/C shRNA transduced B16F10 cells. We found that the transcript levels of LINC complex components such as nesprins 1-4, emerin, as well as of other lamin A/C interactors involved in heterochromatin content and stability (e.g.,Trim28, Tpr or Nup153 (Krull et al., 2010; Kubben et al., 2010)) were all normally transcribed in lamin A/C knockdown B16 cells (Supplementary file 1). Lamin A/C downregulated B16F10 cells exhibited, however, alterations in several hundred genes, including nuclear matrix genes, nuclear body genes, as well as in lamin B receptor (Figure 4D, E and Figure 4-figure supplement 1A-C; Supplementary file 1). Consistent with their similar transmigration properties in vitro and similar capacity to extravasate lung capillaries in vivo, our transcript analysis did not indicate any differences in vascular permeability factors such as VEGF family member levels, MMP levels, or canonical and abundantly expressed cytoskeletal machineries (e.g., Rho GTPases and myosin) (Fig. 4D and Supplementary file 1).

**Figure 4.**
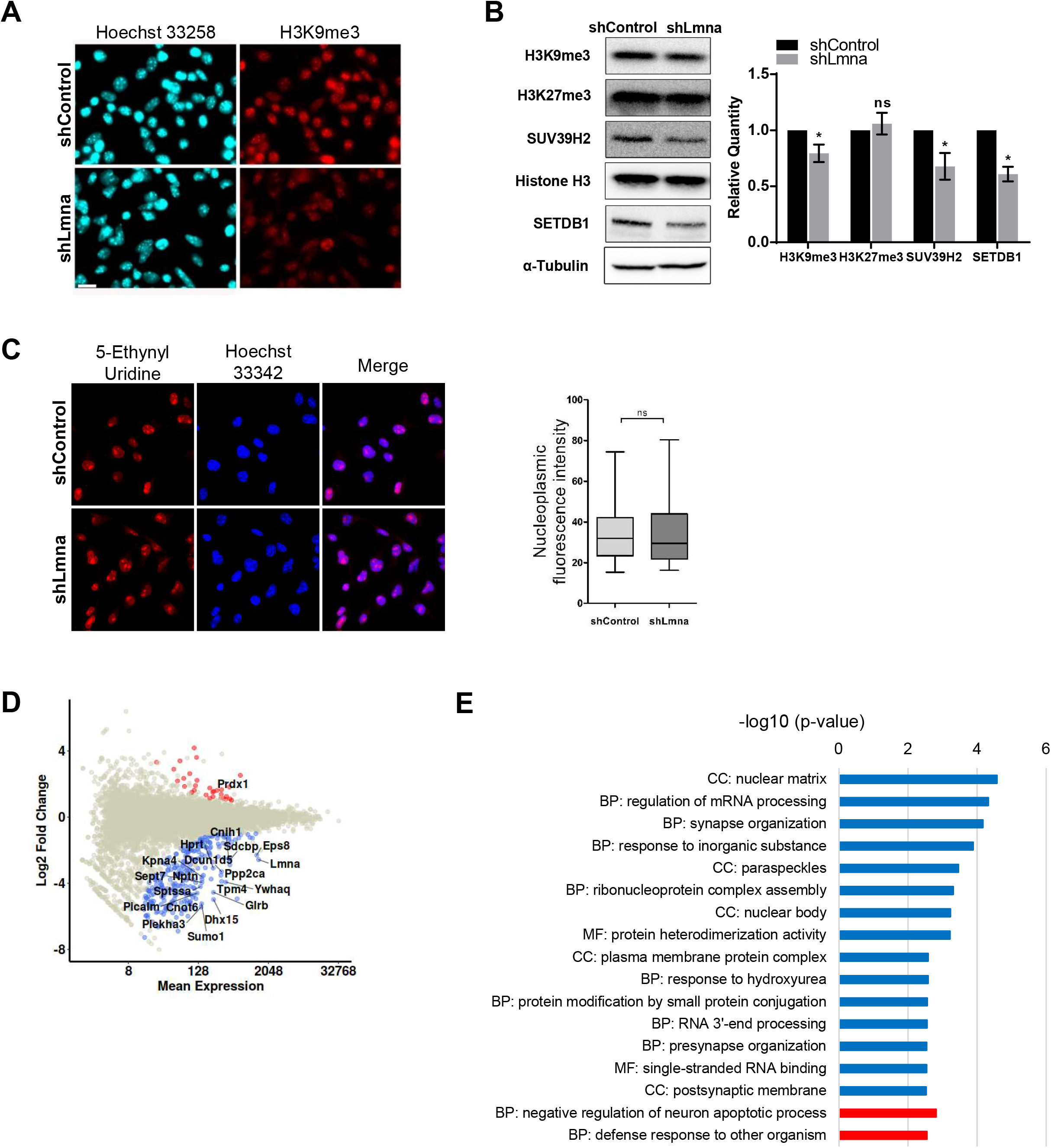
Lamin A/C downregulation reduces heterochromatin content and alters gene transcription. (A) Immunostaining of H3K9me3 (red) in B16F10 shControl and shLmna cells. Hoechst 33258 (cyan) is shown in the left panel. Scale bar, 25 μm. The experiment shown is representative of three. (B) Equal amounts of proteins from B16F10 shControl or shLmna cells, separated by SDS-PAGE and analyzed for the indicated proteins by Western blot analysis. The bar graph represents the mean levels of H3K9me3, H3K27me3 and SUV39H2 normalized to Histone H3 and of SETDB1 normalized to α-Tubulin ± SEM of at least four independent experiments. *p < 0.05. (C) Fluorescence microscopy imaging of 5-ethynyl uridine (EU) incorporation (red) and Hoechst 33342 (blue) in B16F10 shControl and shLmna cells. Cells were grown with 1 mM EU for 1 h. Cells were fixed, permeabilized and treated with Alexa Fluor 594 azide. Nucleoplasmic fluorescence intensity of the EU staining was measured using ImageJ. Data from 3 independent experiments are shown in the boxplot. (D) Differentially downregulated (blue), upregulated (red) and non-differentially expressed genes (grey) are shown (LFC > 1 and adjusted p value < 0.05). The names of the top 20 most significantly expressed genes are indicated in the plot. (E) Gene ontology (GO) enrichment analysis of the top differentially downregulated (blue) and upregulated (red) genes in B16F10 shLmna cells. Biological Process (BP), Molecular Function (MF) and Cellular Component (CC).

### Lamin A/C downregulation does not alter cell proliferation on rigid 2D surfaces but impairs 3D cell growth in spheroids

Our transcriptional analysis did not detect changes in genes involved in melanoma growth (Supplementary file 1). Indeed, melanoma cells knockdown in lamin A/C expression exhibited similar growth rates when seeded at low densities on tissue culture plates (Figure 5A). Interestingly, deficiency in lamin A/C content also did not increase the susceptibility of the nuclei of these melanoma cells to DNA damage: lamin A/C knockdown B16 cells remained as resistant as control melanoma cells to apoptosis and growth arrest induced by the DNA damage and cell cycle arrest inducer etoposide (Dai et al., 2017) (Figure 5B). An additional incubation period without etoposide (Figure 5-figure supplement 1), drove the growth arrested B16 cells into senescence with a slightly higher senescence acquisition exhibited by the lamin A/C knockdown melanoma cells (Figure 5C). Nevertheless, the quantity of R-loops, transient RNA-DNA hybrids which correlate with DNA damage (Crossley, Bocek, & Cimprich, 2019) were comparable for control and lamin A/C knockdown B16 cells (Figure 5-figure supplement 2). These results collectively suggest that lamin A/C knockdown B16F10 cells expand normally in conventional 2D culture conditions, do not exhibit any spontaneous DNA damage, and undergo normal growth arrest induced by a chemical DNA damaging signal.

**Figure 5.**
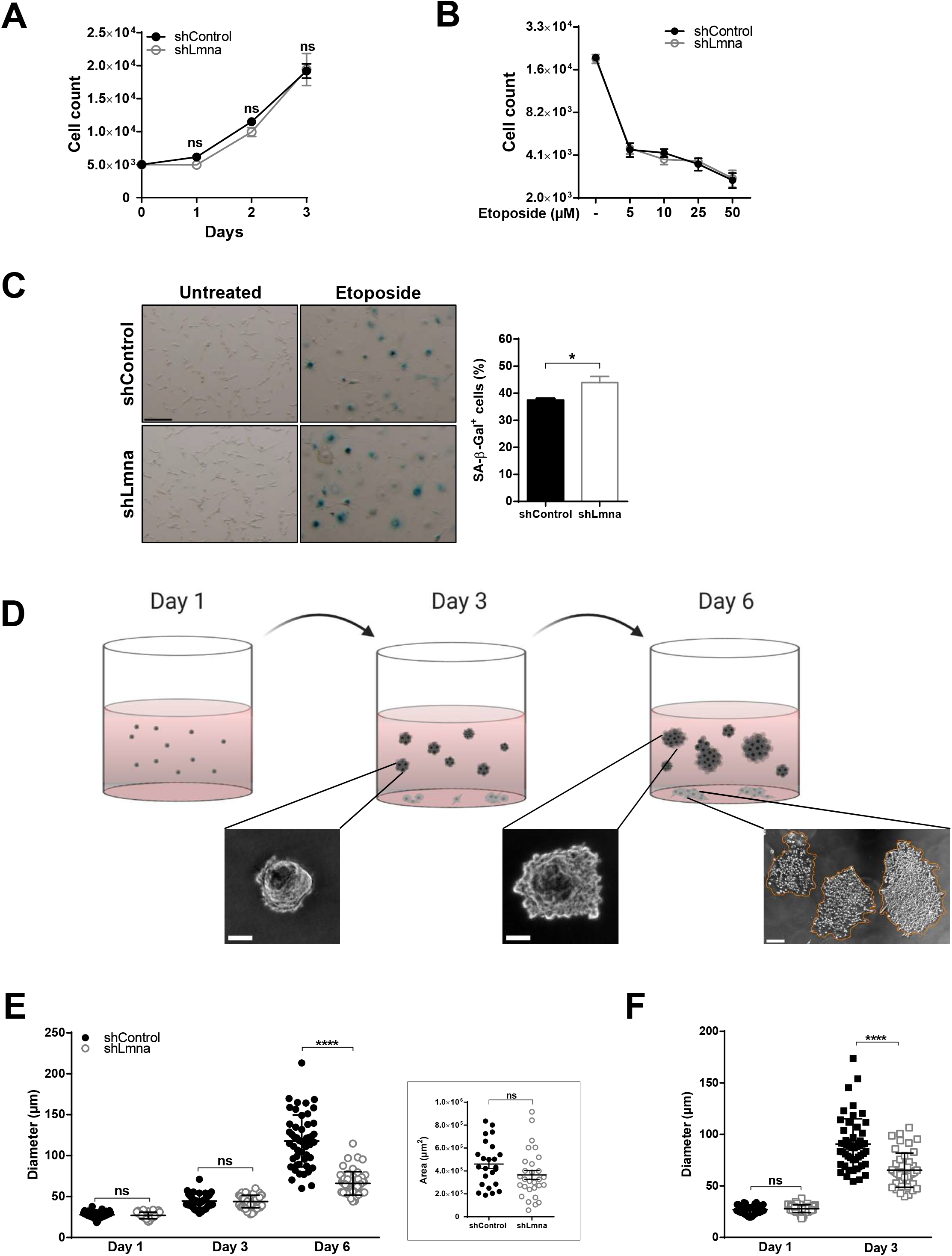
Melanoma proliferation, response to DNA damage, and senescence are insensitive to lamin A/C downregulation. (A) In vitro cell growth of B16F10 shControl and shLmna cells cultured with growth medium changed every 24 h. Values represent the mean ± SEM. The experiment shown is representative of three. (B) Growth arrest induced by etoposide at different concentrations on B16F10 shControl and shLmna cells after treatment for 72 h. Values represent the mean ± SEM. The experiment shown is representative of three. (C) Senescence associated β-galactosidase staining of B16F10 shControl and shLmna cells treated with 5 μM etoposide for 72 h, washed and cultured in regular growth medium for an additional 120 h. The experiment shown is representative of two. Scale bar, 20 μm. *p = 0.0470. (D) Schematic representation of the 3D agarose spheroid assay. Spheroids (dark grey) grown inside agarose while few cells migrate and start dividing at the bottom of the culture dish. At day 6, spheroids were recovered. Representative images of the spheroids derived from B16F10 shControl cells grown in 3D agar (supplemented with 10% FBS) imaged on days 3 and 6. Scale bar, 20 μm. An image of a colony of cells proliferating on the bottom of the soft agar well is depicted at the right. Scale bar, 200 μm. (E) Spheroid diameter, measured on days 1, 3 and 6 after seeding. n = 50 spheroids per group. Values represent the mean ± SEM. The experiment shown is representative of three. ****p < 0.0001. Inset depicts the area of individual cell colonies proliferating on the bottom of the soft agar well, measured on day 6. (F) The diameter of individual spheroids grown for 1 and 3 days in soft agar supplemented with rich growth medium (50% FBS). Values represent the mean ± SEM. The experiment shown is representative of two. ****p < 0.0001.

The soft agar colony formation assay has been a hallmark of cancer survival analysis, as it measures the ability of cells to proliferate in semi-solid matrices (Horibata, Vo, Subramanian, Thompson, & Coonrod, 2015). This readout also allows a direct comparison of cancer cell growth in spheroids vs. in colonies expanded in the same soft agar, while remaining in direct contact with the culture dish 2D surface (Figure. 5D). Interestingly, whereas in the first 3 days both lamin A/C knockdown and control B16F10 cells gave rise to similar sized spheroids (Figure 5E), the ability of lamin A/C high B16 spheroids to further expand during the next 3 days significantly surpassed that of lamin A/C knockdown cells (Figure 5E). In contrast, the growth rate of both B16F10 cell groups settled on the culture plate in the presence of identical soft agar remained similar (Figure 5E, inset). These results were reproduced with a second lamin A/C knockdown B16F10 line expressing a distinct lamin A/C shRNA (Figure 5-figure supplement 3). Remarkably, even in the presence of nutrient rich agar (i.e., a 5-fold higher serum concentration) that accelerated the growth of both lamin A/C high and lamin knockdown B16 cells, the mean size of lamin A/C high B16 cells spheroids was significantly higher than that of lamin A/C low B16 cells (Figure 5F). Interestingly, a negligible fraction of cells isolated from spheroids (9 vs 12 %) underwent apoptosis as depicted by annexin V and propidium iodide staining (Figure 5-figure supplement 4), ruling out the possibility that cells growing at the core of these spheroids have poor accessibility to nutrients. Collectively, our results suggest that whereas initially, lamin A/C knockdown B16F10 normally proliferate under both 2D and 3D conditions, the ability of lamin A/C knockdown cells to expand within 3D spheroids is significantly impaired, possibly due to slower growth of these cells inside large spheroids embedded in 3D environments.

### Lamin A/C deficiency does not affect melanoma growth in distinct primary transplants but results in poorer survival in the lungs and reduced metastasis

In order to validate our in vitro observations, lamin A/C knockdown B16F10 cells or their control counterparts were subcutaneously implanted at 10-20 fold lower numbers than in commonly used orthotopic melanoma models in order to avoid the masking of any potential growth differences. Interestingly, both lamin A/C knockdown and control B16F10 cells grew at similar rates in vivo (Figure 6A). Histological analysis of the orthotopic tumors also failed to show any noticeable differences in the vascular microenvironment (data not shown). Furthermore, both lamin A/C high and low melanoma cells shared similar growth properties when implanted in the mammary fat pad (Figure 6B). Notably, although lamin A/C knockdown B16F10 accumulated at comparable levels to control B16F10 inside recipient lungs within the first 3 days after injection the number of viable lamin A/C knockdown B16F10 recovered from recipient lungs 7 days after injection was significantly reduced relative to control B16F10 cells (Figure 6C). Furthermore, the ability of lamin A/C B16F10 cells to generate metastatic lesions 14 days after injection into recipient mice was dramatically reduced by lamin A/C downregulation (Figure 6D). These results collectively suggest that lamin A/C knockdown B16F10 cells poorly survive in the lungs, in spite of their normal apparent extravasation potential when circulating and entering the blood vasculature.

**Figure 6.**
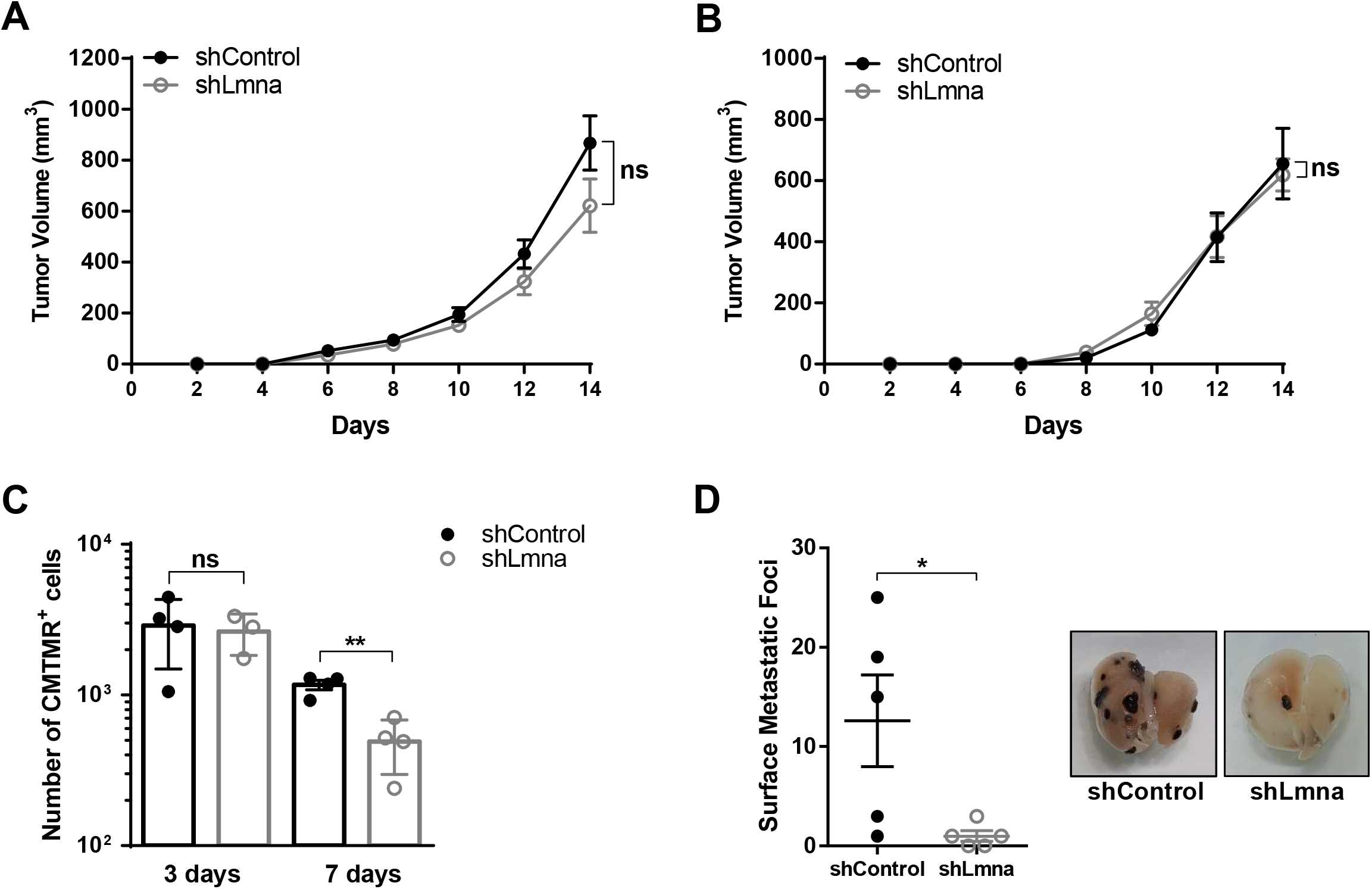
Effects of lamin A/C downregulation on E0771 migration, growth and metastatic potential. (A) 15,000 B16F10 shControl and shLmna cells were implanted subcutaneously in flank or in the mammary fat pad (B) of C57BL/6 mice. Tumor growth was assessed every other day for 14 days post implantation. The experiment shown is representative of three. (C) Number of CMTMR-labeled B16F10 shControl and shLmna present in the lungs of recipient mice, 3 or 7 days after i.v. injection. Data are mean ± SEM. The experiment shown is representative of three. (D) 40,000 B16F10 shControl and shLmna cells were injected in the tail of recipient C57BL/6 mice. After 14 days, animals were euthanized and lungs harvested. Surface metastatic foci were macroscopically counted, results are mean ± SEM, n = 5 for each experimental group. *p = 0.0374. A representative lung image from each group is presented. The experiment shown is representative of three.

### Reduced lamin A/C levels in E0771 breast cancer cells recapitulate the in vitro and in vivo migratory properties and survival deficiencies of lamin A/C knockdown B16 cells

We next reasoned that reduced lamin A/C levels could differentially impact distinct types of cancer cells. We therefore addressed how reduced lamin A/C expression in the bona fide breast cancer cells, the E0771 line, affects their squeezing, migration, epigenetics and growth properties both in vitro and in vivo. Downregulation of lamin A/C levels with conserved lamin B content (Figure 7A), introduced either by shRNA-expressing lentiviral vectors or by siRNA, dramatically facilitated cancer cell squeezing through rigid pores in vitro (Figure 7B and Figure 7-figure supplement 1A, B), in agreement with our findings with B16F10 cells. While the nuclear circularity index of control E0771 cells was higher than that of control B16F10 cells (Figure 7-figure supplement 1C and Figure 2D), the nuclei of these cells underwent significant increases in deformability (i.e. reduced circularity) upon lamin A/C downregulation (Figure 7-figure supplement 1D), reminiscent of the effect of lamin A/C downregulation on B16 nuclei. Nevertheless, the transendothelial migration capacity of these cells in vitro and in vivo was insensitive to reduced lamin A/C expression (Figure 7C and Figure 7-figure supplement 2A-B). Although downregulated lamin A/C expression resulted in reduced H3K9Me3 heterochromatin content and in altered gene expression (Figure 7D and Figure 7-figure supplement 3A-C), it did not increase global RNA transcription rates (Figure 7-figure supplement 4). Furthermore, primary breast cancer growth in vitro or in vivo in the mammary fat pad or in non-orthotopic skin implants was unaffected by lamin A/C downregulation (Figure 7E, F and Figure 7-figure supplement 5). DNA damage-induced growth arrest was also insensitive to downregulated lamin A/C expression (Figure 7G). Nevertheless, and reminiscent of the proliferation results of lamin A/C knockdown B16F10 cells lamin A/C low, E0771 poorly proliferated in spheroids grown in 3D conditions compared to control (Figure 7H). Notably, although Lamin A/C knockdown E0771 cells normally accumulated in recipient lungs when intravenously introduced into syngeneic recipient mice (Figure 7-figure supplement 6). Breast cancer cells with low lamin A/C content failed to generate any metastatic lesions in the lungs (Figure 7I, J). Thus, although reduced nuclear content of lamin A/C does not impair breast cancer growth in the primary tissue and does not alter breast cancer cell extravasation from lung vessels into the lung parenchyma, it dramatically compromises the metastatic potential of these breast cancer cells in this organ, reminiscent of our observations with lamin A/C knockdown melanoma cells.

**Figure 7.**
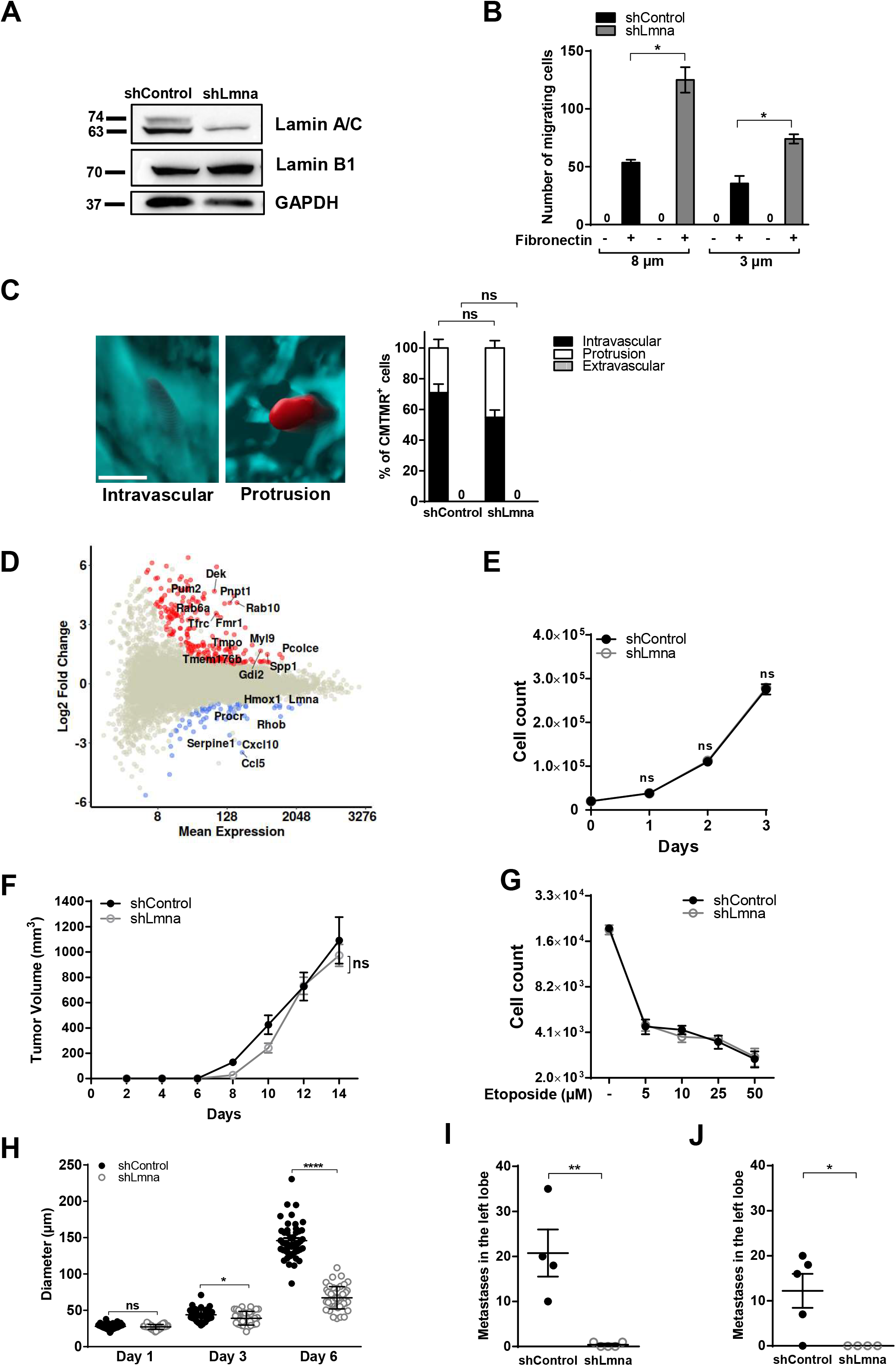
Effects of lamin A/C downregulation on E0771 migration, growth and metastatic potential. (A) Expression levels of lamin A/C and lamin B1 in E0771 cells transduced with control or lamin A/C shRNA. Glyceraldehyde-3-phosphate dehydrogenase (GAPDH) was used as loading control. (B) Haptotactic migration of E0771 shControl and shLmna cells through 8 or 3 μm pores transwell filters in the presence (+) or absence (−) of fibronectin for 4 h. Values represent the mean ± SEM of five fields of view in each experimental group. Results shown are from a representative experiment of three. *p = 0.0240 (8 μm) *p = 0.0361 (3 μm). (C) Representative 3D images of intravascular or protrusive tumor cells across the CD31-labeled lung vasculature together with the percentage of E0771 shControl and shLmna cells present in a volume of 5×10^9^ μm^3^ of the left lung lobe (3 h after injection) counted using Imaris software. The results shown are representative of 3 experiments. Scale bar, 100 μm. (D) Differentially downregulated (blue), upregulated (red) and non-differentially expressed genes (grey) are shown (LFC > 1 and adjusted p value < 0.05). The names of the top 20 most significantly expressed genes are indicated in the plot. (E) In vitro cell growth of E0771 shControl and shLmna cells cultured with growth medium changed every 24 hours. Values represent the mean ± SEM. The experiment shown is representative of two. (F) 10,000 E0771 shControl and shLmna cells were implanted in the mammary fat pad of female C57BL/6 mice. Tumor growth was assessed every other day for 14 days post implantation. The experiment shown is representative of three. (G) Growth arrest induced by etoposide at different concentrations on E0771 shControl and shLmna cells after treatment for 72 hours. Values represent the mean ± SEM. The experiment shown is representative of three. (H) Spheroid diameter, measured on days 1, 3 and 6 after seeding. n = 50 cells per group. Values represent the mean ± SEM. The experiment shown is representative of three. ****p < 0.0001. (I) Experimental lung metastasis of E0771 breast cancer cells. 20,000 E0771 shControl and shLmna cells were intravenously injected in recipient C57BL/6 mice. After 14 days, animals were euthanized and lungs harvested. The number of micrometastases present in the left lung lobe of each lung was determined. The experiment shown is representative of two. **p = 0.0030. (J) 10,000 E0771 shControl and shLmna cells were intravenously injected in recipient C57BL/6 mice. After 30 days, animals were euthanized and lungs harvested. The number of micrometastases present in the left lung lobe of each mouse was determined. The experiment shown is representative of two. *p = 0.0246.

## Discussion

The nucleus is the most bulky organelle in all cells and is protected by a mechanically stable network underlying the inner nuclear membrane termed the nuclear lamina (Swift et al., 2013; Katherine L Wilson & Foisner, 2010; Wolf et al., 2013). The main mechanical obstacle for the extravasation of solid cancer cells across blood vessels in vivo is their stiff nuclei (Shin et al., 2013). The nuclear lamina is thought to affect the shape and the mechanical properties of the nucleus, hence to control cell squeezing through different barriers (Lomakin et al., 2020; Rowat et al., 2013; Wolf et al., 2013; Yadav et al., 2018). The lamina also plays an important role in the maintenance of the nuclear envelope integrity as well as in the organization of the nucleus as a whole (Friedl et al., 2011; Kirby & Lammerding, 2018). The nuclear lamina is also connected to highly condensed chromatin regions (heterochromatin) (Stephens et al., 2012; K. L. Wilson & Berk, 2010) and can affect chromatin conformation, and epigenetics (Fernández-Morera, Calvanese, Rodríguez-Rodero, Menéndez-Torre, & Fraga, 2010; Sullivan et al., 1999). The lamina also controls the entry of key growth control transcriptional factors including the mechanosensitive transcriptional activator YAP (Elosegui-Artola et al., 2017). Lamin A/C expression is reduced in several solid cancers but not others, and so the molecular basis of these changes and their direct link to cancer cell migration, survival and expansion have been under debate due to the complexity and diversity of tumor growth, migration and metastasis (Harada et al., 2014; Kong et al., 2012; Wazir et al., 2013).

We chose to address these key standing questions using two prototypic BL/6 metastatic cell lines, B16F10 melanoma and E0771 breast carcinoma. Taking both in vitro and in vivo reductionist approaches in syngeneic immunocompetent mice, we have systematically assessed how controlled suppression of the two type A lamins affects the specific growth and migratory properties of these cells under distinct physiologically relevant conditions and in different environmental conditions both in vitro and in vivo. Our in vitro findings of enhanced squeezing capabilities of the lamin A knockdown cells in the transwell assay are in full agreement with previous results on the critical role of lamin A/C in the ability of cells of different origins to squeeze through rigid confinements in vitro (Aureille, Belaadi, & Guilluy, 2017; Rowat et al., 2013). However, we found that nuclear deformability has no impact on the overall nuclear squeezing kinetics through endothelial junctions and under endothelial monolayers, indicating that endothelial barriers are highly permissive for nuclear passage and well adapted to accommodate the squeezing of cells with bulky and stiff nuclei. The exceptionally slow rates of tumor transendothelial migration, may also provide the endothelial cytoskeleton with sufficient time to undergo remodeling including activation of contractility machineries to facilitate the squeezing of the relatively stiff nuclei of most tumor cells though junctions or transcellular pores (Khuon et al., 2010; Yadav et al., 2018). In order to validate our in vitro TEM results, we developed a new experimental lung metastasis model to assess in vivo the intrinsic ability of our lamin A/C suppressed tumor cells to squeeze through the lung vasculature. Tumor metastasis into lungs occurs nearly exclusively through the pulmonary capillaries (Miles et al., 2008), an extensive network of relatively impermeable capillaries (Chambers et al., 2002) considered to be poorly permeable compared with vessels targeted by metastatic cells at other organs like the bone marrow and liver (Valastyan & Weinberg, 2011). By introducing a new method to accurately measure the relative efficiency of cancer cell emigration through these vessels, we found that highly invasive cells like melanoma B16 crossed these barriers in vivo independently of their lamin A/C content and irrespectively of their intrinsic nuclear deformability properties. Although poorly invasive, the ability of E0771 cells to cross identical pulmonary vascular barriers was also irrespective of their intrinsic nucleus deformability and lamin A/C content. Our in vivo results were therefore fully consistent with the transmigratory properties of these cells determined in our in vitro setups.

Metastasis involves not only cancer cell extravasation but also intravasation into tumor engulfed blood vessels (Joyce & Pollard, 2009). Although the extravasation capacity of a given cancer cell does not predict its intravasation potential across the tumor associated blood vessels at its original site of dissemination, both our in vitro and in vivo analysis of tumor cell squeezing through different endothelial barriers predict that tumor cell squeezing through endothelial junctions is not affected by lamin A/C downregulation. Furthermore, since the lung capillaries, where the majority of our cancer cell diapedesis took place, are less permeable than tumor associated vessels and were permissive for the squeezing of lamin A/C rich nuclei, it is likely that the leaky vessels surrounding the primary tumors also do not pose a barrier for melanoma or breast cancer intravasation. Likewise, melanoma and breast cancer crossing of lymphatic vessels nearby tumors, considered highly permeable cell barriers (Dyer & Patterson, 2010), is probably insensitive to the lamin A/C content of the tumor nuclei.

Our results argue that the squeezing ability of a given tumor cell through non degradable rigid pores towards a chemotactic or a haptotactic signal does not directly predict the physiological crossing potential of that cell. Although not addressed in our study, it is possible that upon entering the interstitial space, the ability of lamin A/C rich tumor cells to enzymatically degrade glycoprotein and proteoglycan components of collagenous barriers (Bishop, Schuksz, & Esko, 2007) is much more critical for their interstitial motility than the mechanical restrictions imposed by the stiffness of their lamin A/C rich nuclei Indeed, a lamin A/C regulated crosstalk between the nucleus and MT1-MMP has been recently shown to promote a digest-on-demand program in cells squeezing through constricted spaces (Infante et al., 2018). This study strongly suggests that high levels of lamin A/C and optimal connections of these lamins with the cell cytoskeleton (e.g., via the linker of nucleoskeleton and cytoskeleton (LINC) complex) can be in fact essential for directional proteolysis at the cell front.

Our epigenetic analysis revealed lower levels of the constitutive heterochromatin marker H3K9me3 and the methyltransferases that generate it in lamin A/C knockdown B16F10 and E0771 cells, consistent with previous reports on lamin A/C KO MEFs (Liu et al., 2013) and human fibroblasts expressing mutated type A lamins (Scaffidi & Misteli, 2006). Reduced heterochromatin content has been argued to reduce chromatin compaction in B16 melanoma and thereby restrict cell motility (Maizels et al., 2017). The increased squeezing rate of the lamin A/C knockdown B16 cells and E0771 cells that we find in spite of their lower H3K9me3 content sheds new light on these earlier findings. Our data therefore suggest that the altered mechanical properties of our melanoma and breast cancer cell models introduced by lamin A/C downregulation override any inhibitory effects of reduced heterochromatin content on chromatin compaction and cell motility (Maizels et al., 2017). Previous observations showed that depletion of lamin A/C in lung and breast cancer cells, as well as fibrosarcoma cells, significantly increased the likelihood of transient nuclear envelope rupture events and cell death especially when cells were forced to migrate through very tight and rigid barriers (C. M. Denais et al., 2016; Harada et al., 2014). Imaging of HT1080 fibrosarcoma cells invading the collagen-rich mouse dermis in live tumors after orthotopic implantation confirmed that migration-induced nuclear envelope rupture occurs in vivo, particularly during cell division (Vargas, Hatch, Anderson, & Hetzer, 2012) and in individually disseminating cells (C. M. Denais et al., 2016). Nuclear envelope rupture is less prevalent, however, in cells moving as multicellular collective strands (C. M. Denais et al., 2016) or on 2D surfaces (Harada et al., 2014). Notably, both the melanoma and breast cancer cells studied by us grew and survived normally when orthotopically or non orthotopically implanted in distinct organs. Our results therefore argue against faster nuclear rupture of these cells in their primary tissues. On the other hand, lamin A/C knockdown melanoma and breast cancer cells exhibited poor long term survival in the lungs as well as reduced growth inside spheroid assemblies grown in 3D environments, two environments that are far less rigid than tissue culture dishes (Barney et al., 2016; Jaiswal et al., 2017). Therefore, the contribution of lamin A/C to cancer cell growth within spheroids is unlikely to involve classical lamin A/C regulated mechanotransduction which predominates cell-extracellular matrix contacts (Donnaloja, Carnevali, Jacchetti, & Raimondi, 2020).

The contribution of high lamin A/C content to chromatin flexibility and DNA stability is still disputed. On one hand, high lamin A/C content protects the nuclei from rupture and cell apoptosis occurring under extreme mechanical stress such as squeezing through rigid confinements (C. M. Denais et al., 2016; Raab et al., 2016) and also reduces the frequency of spontaneous aneuploidy (Capo-chichi et al., 2011) and DNA damage (Gonzalo, 2014). A-type lamins also participate in the maintenance of telomere homeostasis (Gonzalez-Suarez et al., 2009). Our results extend these findings proposing an additional role of lamin A/C, a protective function of these lamina proteins in multi-cellular assemblies, in the absence of apparent squeezing through rigid confinements. Our findings are the first to implicate lamin A/C as essential protective factors in lung metastasis of multiple types of cancer cells. Our results predict that lamin A/C deficiency impairs cancer cell survival in the lungs to a greater extent than the migratory advantages this deficiency may provide to cancer cells. The relevance of our findings to other types of cancers and metastatic spread in other organs awaits for future studies.

## Materials and methods

### Key resources table

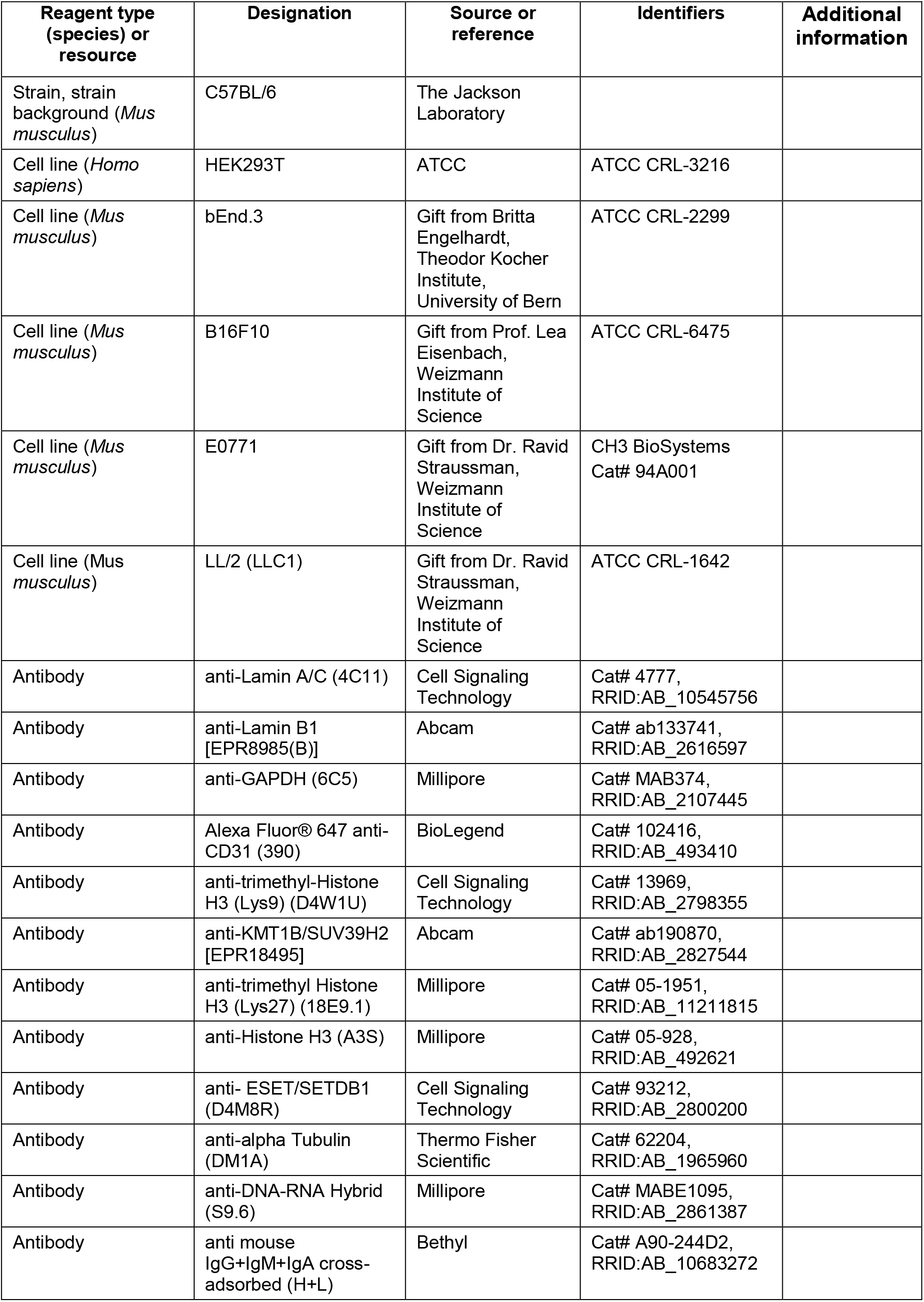
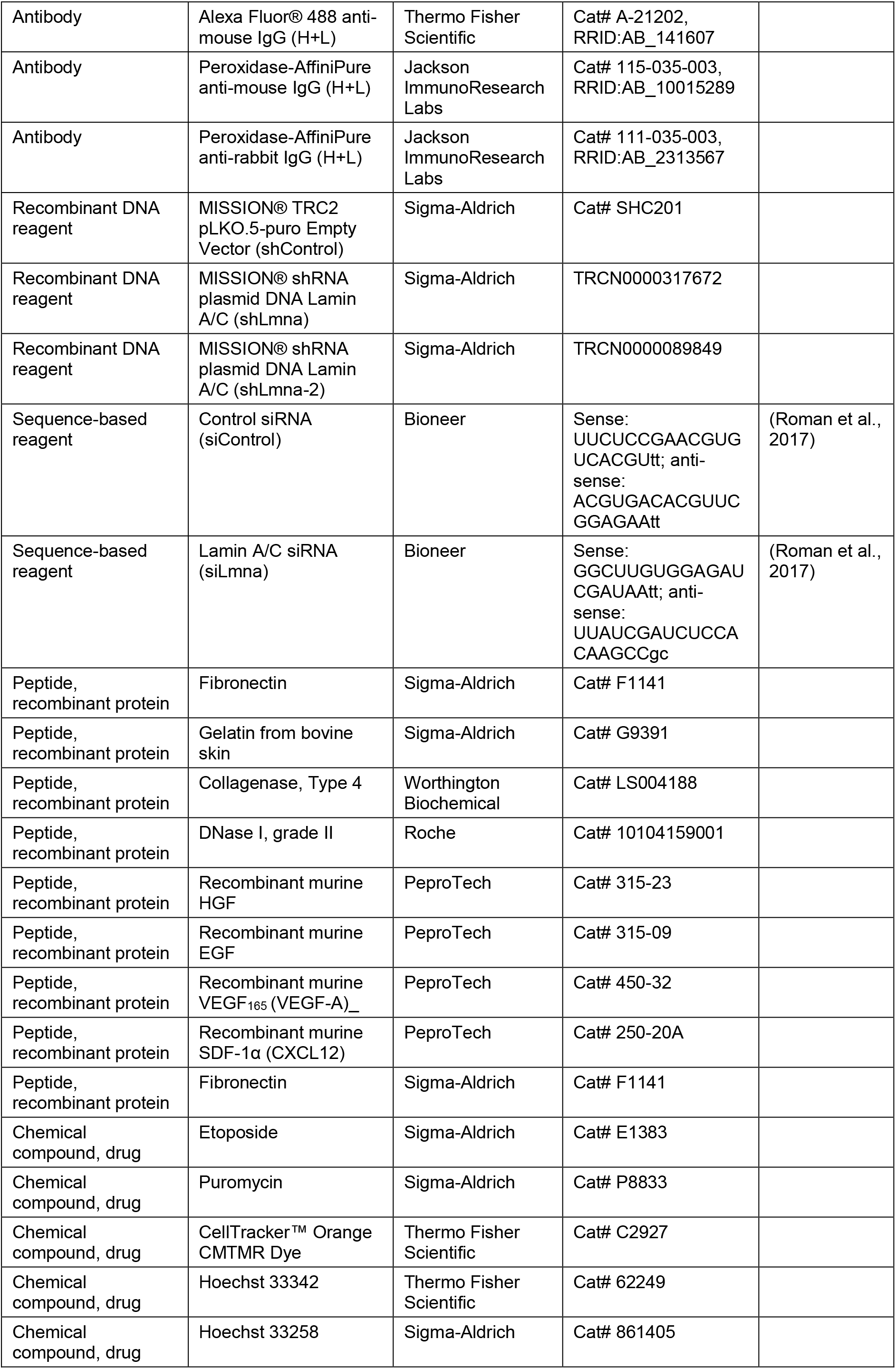
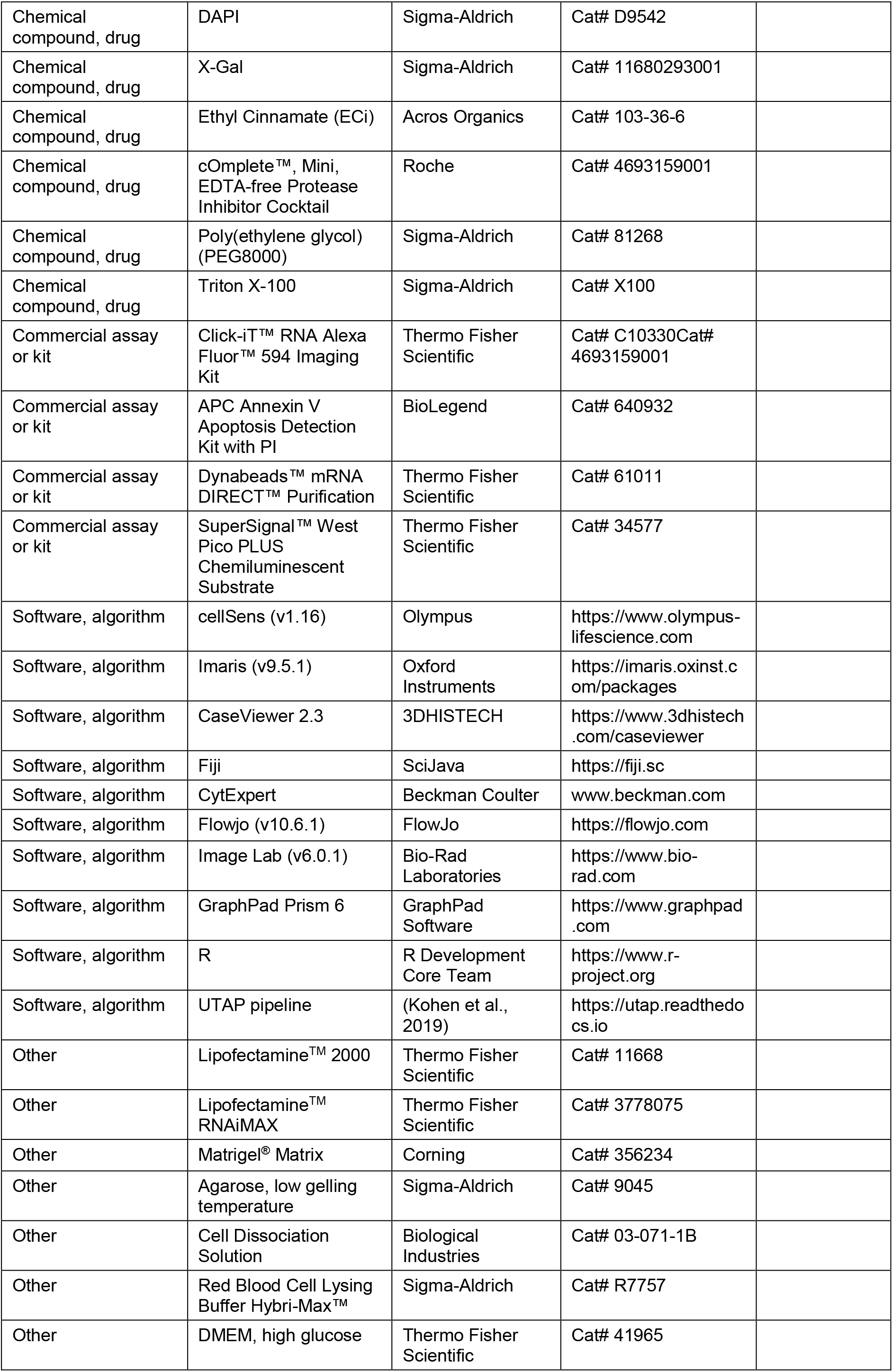

### Cells

Murine melanoma (B16F10) and Lewis Lung Carcinoma (LL/2) cells were grown in DMEM supplemented with 10% FBS. Murine breast adenocarcinoma cells (E0771) were grown in DMEM supplemented with 10% FBS, 1 mM sodium pyruvate and 10 mM HEPES. Human embryonic kidney (HEK293T) and murine brain endothelial (bEnd.3) cells were cultured in DMEM medium supplemented with 10% FBS and 2 mM L-glutamine.

### Mice

Wild-type mice (WT) on C57BL/6 background were maintained in a pathogen-free facility and all animal procedures were approved by the Animal Care and Use Committee of the Weizmann Institute of Science. Male and female 7-to 8-week-old mice were used in all experiments.

### Imaging and Analysis of Tumor Cell Transendothelial Migration

The transmigration assay of tumor cells was performed under shear-free conditions. Murine endothelial bEnd.3 cells (8×10^4^) were seeded in a μ-Slide VI0.4 ibiTreat (ibidi), pre-coated with gelatin (1% in DDW) for 30 min at 37°C. A day later, B16F10 or E0771 cells were labeled with 20 μM Hoechst 33342 for 5 min at 37°C and resuspended in binding medium (Hank’s balanced-salt solution 1X containing 2 mg/ml BSA and 10 mM HEPES, pH 7.4, supplemented with 1 mM CaCl_2_ and 1 mM MgCl_2_) and introduced in the ibidi chamber over a confluent bEnd.3 monolayer. Images were acquired at a rate of one frame every 4-5 min for 4 h using an IX83 inverted microscope (Olympus) equipped with UPlanFLN 20x/0.50 Ph1 ∞/0.17/FN 26.5 objective (Olympus), 49000-ET-DAPI filter set (Chroma). ORCA-Flash4.0LT camera, model: C11440-42U (Hamamatsu). Temperature was maintained at 37°C throughout the assay. For analysis of migratory phenotypes, tumor cells in different fields of view (10-15 cells per field) were individually tracked and categorized using cellSense software 1.16 (Olympus). Close monitoring of individual frames allowed the discrimination of transmigrating tumor cells (TEM) from tumor cells that failed to complete TEM either because of inability to protrude through endothelial junctions (SA) or squeeze their nuclei through these junctions and underneath the endothelial monolayer (SEP). See also Video 2. Notably, we did not observe intercalation of individual tumor cells in between ECs (Reymond et al., 2012). Fiji software was used to determine nuclear circularity of transmigrating tumor cells at different frames of the time-lapse movies. This software was also used to incorporate time codes, labels and scale bars into video segments.

### Tumor Cells Transwell Migration Assay

Fibronectin (1.5 μg/ml in PBS) was coated for 30 min at 37°C onto both sides (for tumor chemotaxis assays) or only on the bottom side (for tumor haptotaxis assays) of 8 or 3 μm hanging cell culture inserts (Millipore, MCEP24H48 and MCSP24H48). After washing the filters with PBS, B16F10 or E0771 cells (4×10^4^) resuspended in DMEM containing 0.1% BSA were introduced into the top chamber. DMEM with 0.1% BSA was inserted in the lower chamber in the presence or absence of the chemoattractant. After 4 or 24 h at 37°C with CO_2_, the cells were fixed with paraformaldehyde (4% in PBS) for 15 min and stained with crystal violet (3% in DDW) for additional 15 min, both at RT. Cells on the upper side of the filter were scraped using a cotton swab whereas cells located on the bottom side were imaged using a SZX16 stereo microscope (Olympus) equipped with SDF PLAPO 1XPF objective (Olympus) set at 10X magnification. DP73 camera (Olympus).

### Generation of Stable shRNA-Expressing Clones

Lentiviruses were produced by co-transfecting HEK293T cells with the shControl or shLmna vectors and three helper plasmids (Gag-Pol, Rev and VSV-G,) using Lipofectamine® 2000. The virus-containing medium was harvested 48 or 72 h after transfection and subsequently precleaned by a brief centrifugation at 600 × g and a 0.45 μm filtration. Viruses were collected and concentrated with a precipitation solution (40% PEG8000 and 2.5N NaCl) as described (Guo et al., 2012) and stored at −20°C overnight. A day later, the medium was thawed and centrifuged at 2,400 × g for 30 min at RT. The viral pseudoparticles were resuspended in 200 μl culture medium and mixed with 2.5×10^4^ of tumor cells for 12 h. 36 h after viral infection, puromycin was added to the culture medium at a concentration of 2.5 μg/ml (B16F10) or 3 μg/ml (E0771) and tumor cells were selected and expanded, replacing growth medium every 48 h. The mean knockdown levels were assessed by Western blotting.

### Transient siRNA transfection

B16F10 cells (1×10^5^) were transfected with siRNA (20 nM) using Lipofectamine™ RNAiMAX (Thermo Fisher Scientific) following the manufacturer’s instructions. 72 h after transfection tumor cells were either lysed to quantify the mean knockdown protein levels or utilized in experimental assays. Lamin A/C siRNA (siLmna) sequence used in this study was described in (Roman et al., 2017).

### Western Blotting

Cells were grown as described above, washed twice with ice-cold PBS, scraped into lysis buffer (25 mM Tris pH 7.5, 1 mM EDTA, 0.5 mM EGTA, 150 mM NaCl, 1% NP-40, 0.2% SDS, 2 mM Na_3_VO_4_, 1 mM NaF, 10 mM Nappi, 80 mM b-glycerol phosphate and a protease inhibitor tablet), and kept on ice for 30 min with occasional vortexing. Thereafter, lysates were centrifuged at 14,000 × g for 15 min at 4°C. The supernatant was collected and protein concentration was determined by BCA protein assay (Thermo Fisher Scientific). The protein suspension was separated by gel electrophoresis followed by transfer to nitrocellulose membranes, and blocking with non-fat milk (5% in PBS-T) for 1 h at RT. Immunoblotting was performed overnight at 4°C according to the manufacturers’ guidelines. Antibody binding to membrane blots was detected using horseradish peroxidase conjugated secondary antibodies for 1 h at RT, followed by development with a chemiluminescence substrate (Thermo Fisher Scientific). Chemiluminescence was detected using the ChemiDoc MP (Bio-Rad Laboratories) imaging system.

### Immunofluorescence Staining

B16F10 or E0771 cells (1.5×10^4^) were seeded into a μ-Slide VI0.4 ibiTreat (ibidi) pre-coated with fibronectin (10 μg/ml in PBS) for 30 min at 37°C. The next day cells were rinsed with ice-cold PBS and fixed with paraformaldehyde (4% in PBS) for 15 min at RT followed by permeabilization with Triton X-100 (0.25% in PBS) for 15 min at RT and blocking with goat serum (10% in PBS) for 20 min at 37°C. The cells were then incubated with anti-lamin A/C antibody (1:100) for 1 hour at RT, washed with ice-cold PBS three times and incubated with an Alexa Fluor 488 conjugated secondary antibody (1:200) for 1 h at RT. Cells were imaged using an IX83 inverted microscope (described above) equipped with an UPlanFLN 40X 0.75 Ph2 ∞/0.17/FN 26.5 objective (Olympus), 49002-ET-EGFP (FITC/Cy2) filter set (Chroma). Alternatively, cells were fixed with paraformaldehyde (3% in PBS) at RT for 5 min followed by fixation in methanol at −20°C for another 5 min. H3K9me3 was detected with rabbit monoclonal anti H3K9me3 and DNA was stained with Hoechst 33258. Immunostaining images were collected using an Olympus IX81 fluorescent microscope equipped with a coolSNAP HQ2 CCD camera (Photometrics).

### Imaging of Tumor Cell Nuclear Dynamics

B16F10 or E0771 cells (2×10^4^) were trypsinized, labeled in suspension with 20 μM Hoechst 33342, resuspended in binding medium (composition described above), and introduced in a μ-Slide VI0.4 ibiTreat (ibidi) over a bEnd.3-deposited basement membrane extracellular matrix. Images were acquired at a rate of one frame every 4-5 min for 2 h using an IX83 Inverted Microscope (described above). Temperature was kept at 37°C throughout the duration of the assay. Background was subtracted for the fluorescent channel using cellSense 1.16 (Olympus) software. FiJi (SciJava) software was used for title and time code labeling and determination of the nuclear circularity.

### Light Sheet Fluorescent Microscopy of Tumor Cells and Lung Vasculature

B16F10 (2×10^4^) or E0771 (10^4^) cells labeled with CMTMR dye (Thermo Fisher Scientific), 10 μM for 30 min according to the manufacturer’s instructions, were injected in the retro-orbital sinus of recipient mice. Euthanasia by administration of sodium pentobarbital (200 mg/Kg) was practiced 3 hours later. Blood capillaries were labeled 15 min before the animal sacrifice by intravenous injection of 6 μg of an Alexa 647-conjugated anti-CD31 mAb. Immediately after the sacrifice, mice were transcardially perfused with PBS and the lungs inflated via the trachea with low gelling agarose (Sigma-Aldrich), subsequently fixed with paraformaldehyde (4% in PBS) for 2 h, dehydrated and cleared using ethyl cinnamate as described in (Klingberg et al., 2017). Cleared intact lung lobes, were imaged using an Ultramicroscope II (LaVision BioTec) operated by the ImspectorPro software (LaVision BioTec). For excitation light sheet was generated by a Superk Super-continuum white light laser (emission 460 nm – 800 nm, 1 mW/nm – 3 (NKT photonics), followed by specific excitation filters per channel. For detection optics microscope was equipped with a single lens configuration - 4X objective - LVBT 4X UM2-BG, with an adjustable refractive index collar set to the RI of 1.56. Images were acquired by an Andor Neo sCMOS camera (2,560 × 2,160, pixel size 6.5 μm × 6.5 μm, Andor). Z stacks were acquired in 3 μm steps. Channel configuration for GFP and EGFP excitation 470\40 emission 525\50, for CMTMR, excitation 560\40 emission 630\75, and for CD31-AF647 excitation 640\30 emission 690\50.

### Image Reconstruction and Analysis

Three-dimensional rendering of LSFM was performed via Imaris software (Oxford Instruments). Surfaces of CMTMR-labeled tumor cells were created using volume (comprised between 1000 and 25000 μm^3^) and intensity (max of red fluorescent channel) as defining features to unequivocally separate them from background signals. Each cell was individually segmented and its distance was measured with respect to the CD31-labeled blood vessels: intravascular, extravascular or protruding. See also Videos 6-8.

### Determination of Tumor cell Accumulation in Washed Lungs

B16F10 (2×10^4^) or E0771 (10^4^) cells labeled with 10 μM CMTMR for 30 min according to the manufacturer’s instructions, were resuspended in PBS and injected into the retro-orbital sinus of recipient mice. Euthanasia by administration of sodium pentobarbital (200 mg/Kg) was practiced 3 or 72 h later. Immediately thereafter, mice were transcardially perfused with PBS and the lungs were extracted, minced and incubated in RPMI-1640 containing collagenase type 4 (1.5 mg/ml) and DNase I (20 μg/ml) at 37°C for 45 min. Lung cell suspensions were pushed through a 100 μm cell strainer and centrifuged at 0.2 × g or 5 min at 4°C. RBCs were subsequently lysed with an RBC lysis buffer (Sigma Aldrich). The cells were resuspended in ice-cold FACS buffer (PBS with 1% BSA, 0.1% sodium azide and 5 mM EDTA), filtered through a 70 μm strainer and analyzed using a CytoFLEX flow cytometer (Beckman Coulter).

### 5-Ethynyl Uridine (EU) Labeling of Cultured Cells

B16F10 or E0771 (either shControl or shLmna expressing) cells were grown on serum coated glass coverslips. 1 mM EU was added to the culture medium and cells were kept for 1 h at 37°C in 5% CO_2_. Cells were then washed with PBS and fixed with 3.7% formaldehyde for 15 min at RT. The fixative was removed and cells were washed twice with PBS, followed by permeabilization with 0.5% Triton X-100 for 15 min at RT. Cells were washed twice with PBS and incorporated EU was detected by click chemistry using a fluorescent azide following the manufacturer’s guidelines of Click-iT RNA Imaging Kit (Thermo Fisher Scientific). Following the Click-iT reaction (30 min at RT in the dark), cells were rinsed twice with a rinse buffer. Nuclear staining was performed with Hoechst 33342. Cells were imaged using a Zeiss LSM 710 point scanning confocal microscope with stacking acquisition and generation of maximum intensity projection images. Nucleoplasmic fluorescence intensity measurements were performed in Fiji, and statistical significance was assessed through Mann-Whitney U, two-tailed, test.

### Bulk MARS-seq protocol and sequencing

RNA was isolated from 10,000 cells from each cell line using Dynabeads^®^ mRNA Direct Kit (Thermo Fisher Scientific). Libraries for RNA-seq were prepared using a modified version of TranSeq, as described (Jaitin et al., 2014). Briefly, RNA was reversed transcribed with MARS-seq barcoded RT primers in a 10 μl volume with the Affinity Script kit (Agilent). Reverse transcription was analyzed by qRT-PCR and samples with a similar CT were pooled (up to eight samples per pool). Each pool was treated with Exonuclease I (NEB) for 30 min at 37°C and cleaned by 1.2× volumes of SPRI beads (Beckman Coulter). Next, the cDNA was converted to double-stranded DNA with a second strand synthesis kit (NEB) in a 20 ml reaction, incubating for 2 h at 16 °C. The product was purified with 1.4× volumes of SPRI beads, eluted in 8 μl and in vitro transcribed (with the beads) at 37 °C overnight for linear amplification using the T7 High Yield RNA polymerase IVT kit (NEB). Following IVT, the DNA template was removed with Turbo DNase I (Ambion) 15 min at 37 °C and the amplified RNA (aRNA) purified with 1.2 volumes of SPRI beads. The aRNA was fragmented by incubating 3 min at 70 °C in Zn^2+^ RNA fragmentation reagents (Ambion) and purified with 2× volumes of SPRI beads. The aRNA was ligated to the MARS-seq ligation adapter with T4 RNA Ligase I (NEB). The reaction was incubated at 22 °C for 2 h. After 1.5× SPRI cleanup, the ligated product was reverse transcribed using Affinity Script RT enzyme (Agilent) and a primer complementary to the ligated adapter. The reaction was incubated for 2 min at 42°C, 45 min at 50°C, and 5 min at 85°C. The cDNA was purified with 1.5× volumes of SPRI beads. The library was completed and amplified through a nested PCR reaction with 0.5 mM of P5_Rd1 and P7_Rd2 primers and PCR ready mix (Kappa Biosystems). The amplified pooled library was purified with 0.7× volumes of SPRI beads to remove primer leftovers. Library concentration was measured with a Qubit fluorometer (Life Technologies) and mean molecule size was determined with a 2200 TapeStation instrument. RNA-seq libraries were sequenced using the Illumina NextSeq® 500 High Output v2 Kit (75 cycles).

### Bioinformatics analysis

Samples were demultiplexed using the barcode present in the R2 read. The analysis was performed using the UTAP pipeline (Kohen et al., 2019). In brief, UMI sequences present in the R2 read were inserted in the read name of R1 sequence file using a python script. Cutadapt was used to trim low quality, poly A and adapter sequences (Martin, 2011), (parameters: -a AGATCGGAAGAGCACACGTCTGAACTCCAGTCAC -a “A{10}” –times 2 -u 3 -u -3 -q 20 -m 25). Sequences were mapped to the UCSC mm10 mouse genome using STAR (Dobin et al., 2013) v2.4.2a (parameters: – alignEndsType EndToEnd, –outFilterMismatchNoverLmax 0.05, –twopassMode Basic, –alignSoftClipAtReferenceEnds No). The pipeline quantified the 3’ of RefSeq annotated genes (1,000 bases upstream of the 3’ end and 100 bases downstream) using HTSeq count (Anders, Pyl, & Huber, 2015) and a modified Refseq gtf file (downloaded from igenomes UCSC). UMI information was integrated into the BAM files as tags, using a python script. UMI counting was performed after marking duplicates (in-house script) using a modified HTSeq-count. DESeq2 (Love, Huber, & Anders, 2014) was used for normalization and detection of differentially expressed genes. Raw p-values were adjusted for multiple testing using the procedure of Benjamini and Hochberg. Genes were considered to be differentially expressed if their mean normalized expression was greater than 5, the absolute value of the log2FoldChange was greater than 1, and the adjusted p-value was less than 0.05. Batch effects were removed from the read counts using the ComBat function from the sva R package (Johnson, Li, & Rabinovic, 2007). The normalized, batch corrected and log2 transformed read counts outputted by ComBat were used to draw plots. K-means clustering of differentially expressed genes was performed using the Partek® Genomics Suite® software, version 6.6 (Partek Inc., St. Louis, MO, USA). Functional analysis of the differentially expressed genes was performed using https://metascape.org/gp/index.html#/main/step1, Metascape (Zhou et al., 2019). Significantly enriched GO terms for up and downregulated genes were extracted from this analysis. Sequencing data have been deposited in NCBI’s Gene Expression Omnibus and are accessible through GEO Series accession number GSE146730.

### Cell Growth on 2D Culture Plates

B16F10 or E0771 cells were seeded at a low density of 5×10^3^/well in a 6-well plate. Puromycin-containing growth medium was replaced every 24 h throughout the duration of the assay (72 h). To determine the number of viable cells proliferating on the plates, cells were trypsinized and counted by flow cytometer every 24 h.

### Cell Growth in 3D Soft Agar

B16F10 or E0771 cells were suspended at a density of 4×10^3^/ml of 0.3% low gelling agarose in DMEM +10% FBS and equilibrated at 4°C for 15 min in 12 well plate as described (Horibata et al., 2015). The cell agar suspension was warmed to 37°C and cultured in a humidified incubator containing 5% CO_2_. Spheroid formation was monitored on the first, third and 6th day using a IX83 inverted microscope (Olympus) equipped with UPlanFLN 4x/0.13 Ph1 ∞/-/FN 26.5 objective (Olympus). In parallel, cancer cell colonies grown directly on the bottom of each well were also images. To isolate individual cells from the spheroids grown in 3D conditions, the layer of agarose containing the spheroids was mechanically removed and solubilized in PBS pre-warmed at 45°C. Spheroids were spun down at 0,8 × g for 5 min at RT, trypsinized with warm trypsin B to recover single cells and stained with annexin V and propidium iodide following the manufacturer instructions.

### Primary Tumor Growth in vivo

A suspension of 1.5×10^4^ B16F10 or 10^4^ E0771 (either shControl or shLmna expressing) cells in 50 μl of Matrigel^®^ Matrix mixed with PBS (at 1:1 v/v), was inoculated subcutaneously in the flank (B16F10) or in the mammary fat pad (E0771) of recipient mice. Tumor size (volume) was assessed throughout the duration of the experiment, by vernier caliper measurements of length (L) and width (W) and the tumor volume (V) was calculated using the formula: V = (L × W × W)/2. After 14 days, the animals were euthanized by CO_2_ inhalation and the tumor was extracted, weighted and fixed in paraformaldehyde (4% in PBS).

### Experimental Lung Metastases

B16F10 (4×10^4^) or E0771 (10^4^ or 2×10^4^) either shControl or shLmna expressing cells were suspended in 200 μl PBS + 0.25 mM EDTA and injected into the tail vein of recipient mice. Animals were euthanized by administration of sodium pentobarbital (200 mg/Kg) 14 or 28 days later. Immediately after the sacrifice, mice were transcardially perfused with PBS and the lungs were extracted and visually analyzed for the presence of surface metastatic foci, subsequently stored in 4% PFA for 24 h and 1% PFA at 4°C for long term storage. Paraffin embedding and H&E staining of 5 μm-thick sections were performed by the histology core unit of the Weizmann Institute of Science. Sections were digitalized using a Pannoramic SCAN II (3DHISTECH) and analyzed using CaseViewer software (3DHISTECH).

### Senescence-Associated-β-Galactosidase (SA-β-Gal) Activity

For SA-β-gal activity assay, B16F10 cells (shControl or shLmna) were treated with 5 μM etoposide for 72 h, washed and left in regular culture medium (containing 2.5 μg/ml of puromycin) for an additional 120 h. Cells were then fixed with 0.5% glutaraldehyde in PBS for 15 min, washed twice with PBS supplemented with 1 mM MgCl_2_ (pH 5.5) and stained with PBS containing 1 mM MgCl_2_ (pH 5.5) and supplemented with 1 mg/ml X-gal, 5 mM K_3_Fe[CN]_6_ and 5 mM K_4_Fe[CN]_6_ for 5 h at 37°C in the dark, as described (Krizhanovsky et al., 2008). Cells were washed with warm PBS, fixed with 4% paraformaldehyde and imaged using aSZX16 stereo microscope (Olympus) equipped with SDF PLAPO 1XPF objective (Olympus) set at 10X magnification. DP73 camera (Olympus).

### R-loops quantification

B16F10 cells were grown on sterile coverslips and R-loops were detected with the S9.6 antibody following cell fixation and permeabilization with 100% ice-cold methanol and acetone for 10 min and 1 min on ice, respectively. Incubation with primary antibodies was followed by incubation with Dy488-secondary antibody (Bethyl). Nuclei were stained using DAPI and coverslips were assembled in Vectashield Mounting Medium (H-1000, Vector Laboratories). All the washing steps were done with PBS containing 0.05% (v/v) Tween 20. Images were acquired using Confocal Laser Point-Scanning Microscope Zeiss LSM 710. Fluorescence intensity of the nucleoplasmic staining of 90 cells from three independent experiments was assessed for each condition, and statistical significance was assessed through Mann-Whitney U, two-tailed, test.

### Statistical analysis

Data in graphs are represented as means ± SEM. All tests of statistical significance were performed using a two-tailed Student’s t-test with GraphPad Prism software, with the exception of analysis of Figure 6C and Figure 5-figure supplement 1 for which Mann-Whitney U, two-tailed, test was used. Significance was set to p <0.05. Statistical details of experiments can be found in the figure legends. For figures 6D and 7D, raw p-values were adjusted for multiple testing using the procedure of Benjamini and Hochberg while genes were considered to be differentially expressed if their mean normalized expression was greater than 5, the absolute value of the log2FoldChange was greater than 1, and the adjusted p-value was less than 0.05.

## Acknowledgements

We thank Edgar Gomes (Instituto de Medicina Molecular, Lisbon, Portugal), Valery Krizhanovsky and Avri Ben-Ze’Ev (Weizmann Institute) for providing reagents and for helpful discussions. The RNA-Seq work was performed under the guidance of Drs. Merav Kedmi and Hadas Keren-Shaul from the Sandbox unit of the Life Science Core Facility of Weizmann Institute of Science. R.A. is the Incumbent of the Linda Jacobs Chair in Immune and Stem Cell Research. His research is supported by the Israel Science Foundation (grant no. 791/17), the Minerva Foundation, Germany, GIF (grant number I-1470-412.13/2018), as well as grants from the Moross Integrated Cancer Center, Yeda-Sela Center for Basic Research, Helen and Martin Kimmel Institute for Stem Cell Research, the Meyer Henri Cancer Endowment, and from William and Marika Glied and Carol A. Milett. S.N and S.M. are funded by the Wellcome Trust (grant 098291/Z/12/Z to S.N.). G.G’s research is supported by an Israel Cancer Association grant no. 20201181.

## Author Contributions

Francesco Roncato, Methodology, Validation, Formal analysis, Investigation, Data curation, Writing - original draft, Writing - review and editing, Visualization; Ofer Regev, Methodology, Investigation; Sara W. Feigelson, Validation, Investigation; Writing - original draft, Writing - review and editing, Supervision; Sandeep Kumar Yadav, Investigation; Lukasz Kaczmarczyk, Validation, Investigation, Formal Analysis, Visualization; Nehora Levi, Validation, Investigation, Formal Analysis, Visualization; Diana Drago-Garcia, Software, Formal Analysis, Visualization; Samuel Ovadia, Investigation; Marina Kizner, Investigation; Yoseph Addadi, Supervision, Writing-original draft; João C. Sabino, Validation, Investigation, Formal Analysis, Visualization; Yossi Ovadya, Resources, Investigation; Sérgio F. de Almeida, Resources, Supervision, Writing-original draft; Ester Feldmesser, Software, Formal analysis, Visualization; Gabi Gerlitz, Resources, Supervision, Writing-original draft; Ronen Alon, Conceptualization, Writing - original draft, Writing - review & editing, Supervision, Project administration, Funding acquisition.

## Video Legends

**Video 1**

**Transendothelial migration of B16F10 murine melanoma**

Time-lapse movie of an Hoechst-labeled B16F10 crossing a bEnd.3 endothelial monolayer. The contours of the tumor cell leading edges and nucleus are outlined in each image in yellow and red respectively. Elapsed time is designated as h:mm:ss. Scale bar, 20 μm

**Video 2**

**Migratory phenotypes of B16F10 cells over a bEnd.3 endothelium**

Time-lapse movie divided in four quadrants in which different B16F10 cells display a unique phenotype when interacting with a bEnd.3 endothelial monolayer. Represented clockwise there are Round, Spread Above (SA), subendothelial pseudopodium (SEP) and transendothelial migration (TEM) phenotypes. Elapsed time is designated as h:mm:ss. Scale bar, 20 μm

**Video 3**

**Transendothelial migration of B16F10 shControl vs shLmna cells**

Time-lapse movie depicting a Hoechst labeled B16F10 cell shControl (left) and shLmna (right) crossing a bEnd.3 endothelium. The contours of the tumor cell leading edges and nucleus are outlined in yellow and red respectively. Elapsed time is designated as h:mm:ss. Scale bar, 20 μm.

**Video 4**

**Nuclear deformability of B16F10 shControl vs shLmna cells**

Time-lapse movie depicting the Hoechst-labeled nuclei (green) of B16F10 shControl (left) and shLmna (right) cells interacting with a bEnd.3-deposited basement membrane. Elapsed time is designated as h:mm:ss. Scale bar, 20 μm.

**Video 5**

**Light sheet microscopy of tumor cells, bronchial structures and lung vasculature**

Three-dimensional animated visualization of a section of murine lung lobe. CMTMR-labeled tumor cells (red) and autofluorescent bronchial structures (green) can be observed from seconds 0 to 21 (movie length). CD31-labeled lung vasculature (cyan) can be observed from seconds 22-40 (movie length).

**Video 6**

**Example of an intravascular B16F10 cell**

Three-dimensional animated visualization of a CMTMR-labeled B16F10 cell (red) located inside a CD31-labeled lung vasculature (cyan). Scale bar, 100 μm.

**Video 7**

**Example of a protruding B16F10 cell**

Three-dimensional animated visualization of a CMTMR-labeled B16F10 cell (red) protruding through the CD31-labeled lung vasculature (cyan). Scale bar, 100 μm

**Video 8**

**Example of an extravascular B16F10 cell**

Three-dimensional animated visualization of a CMTMR-labeled B16F10 cell (red) located outside the CD31-labeled lung vasculature (cyan). Scale bar, 100 μm

**Supplementary File 1**

DeSeq2 analysis output and gene expression levels in enriched Gene Ontology terms.

---

**Figure 1–figure supplement 1.**
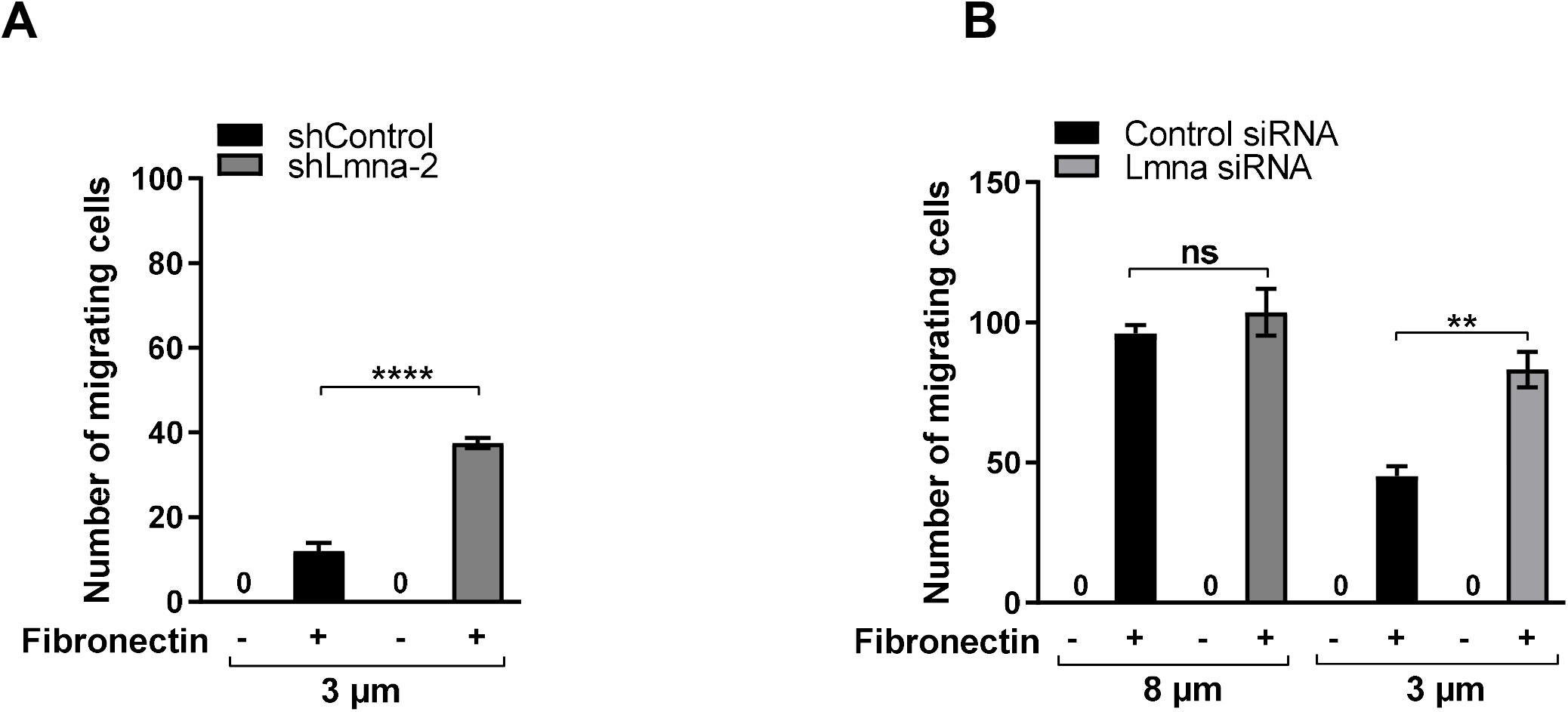
Lamin A/C deficiency increases B16 melanoma cell chemotaxis through rigid pores. (A) Haptotactic of B16F10 shControl vs shLmna-2 through 3 μm pores transwell filters coated (+) or uncoated (−) with fibronectin (1.5 μg/ml) for 4 h. **** p < 0.0001. (B) Migration of B16F10 cells towards HGF (50 ng/ml) measured through 8 or 3 μm pore transwell filters after 4 h. Data are mean ± SEM. The results shown are representative of 2 independent experiments. **p = 0.0076.

**Figure 3–figure supplement 1.**
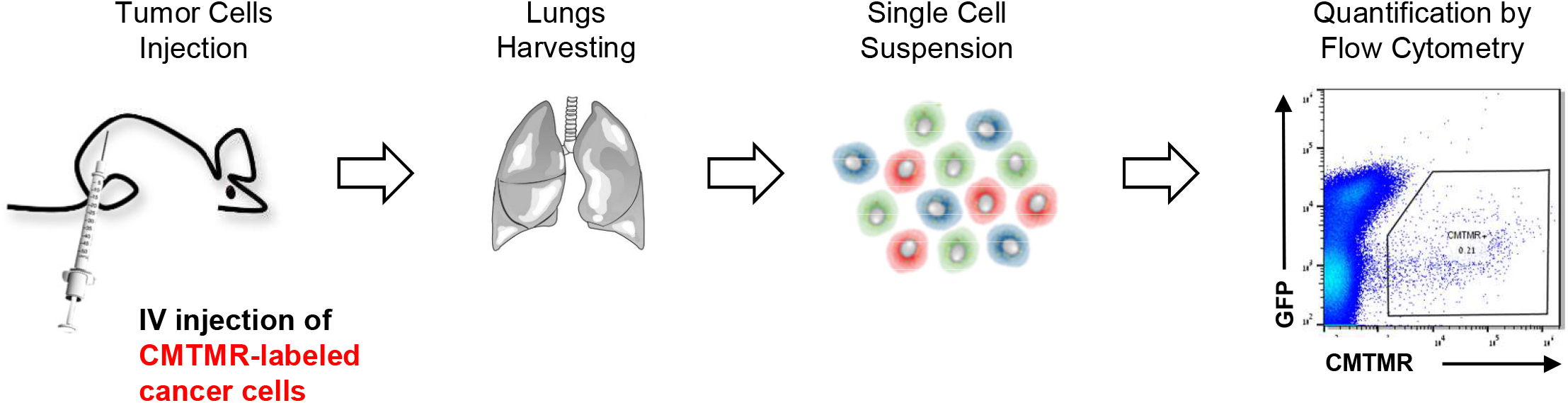
Quantification of cancer cells accumulation in lungs. Scheme depicting the experimental pipeline necessary to quantify single cell accumulation in lungs of recipient mice. Cancer cells were labeled with CMTMR orange cell tracker for 30 min, washed and i.v. injected into recipient WT mice. After 3 hours, 3 or 7 days the mice were euthanized, their lungs harvested, minced and digested into single cell suspension and subsequently analyzed by flow cytometry.

**Figure 4–figure supplement 1.**
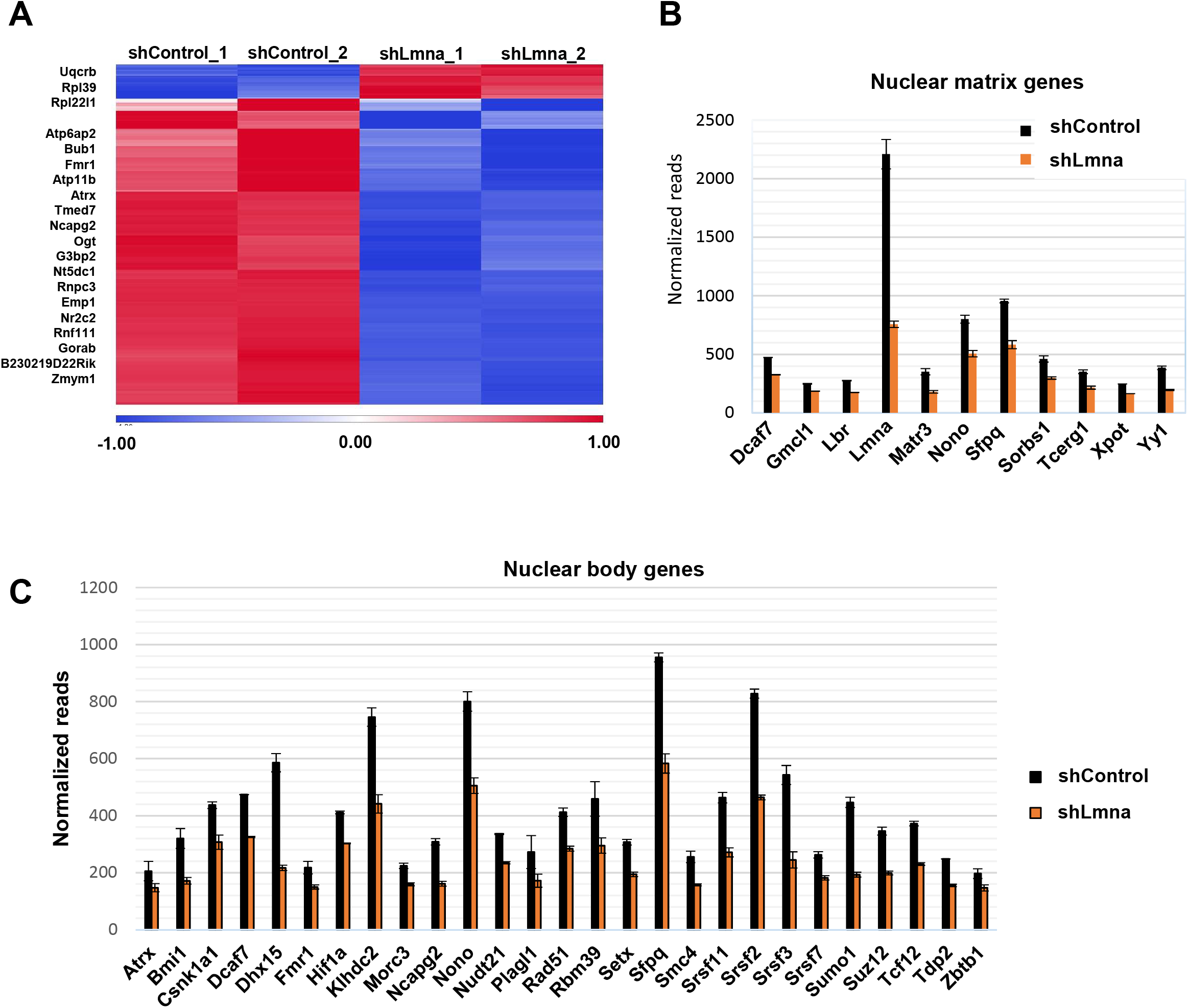
Lamin A/C downregulation alters gene transcription. (A) Heatmap based on clustering of the 300 differentially expressed genes. Red and blue bars represent positive and negative changes, respectively, and the intensity of the color represents the standardized log2 expression level. The top differentially expressed (DE) genes are shown on the left. The full list is available in Table S1. (B, C) Individual gene expression levels of genes belonging to the enriched gene ontology (GO) terms: nuclear matrix (B) and nuclear body (C), summarized in Fig. 6E.

**Figure 5–figure supplement 1.**
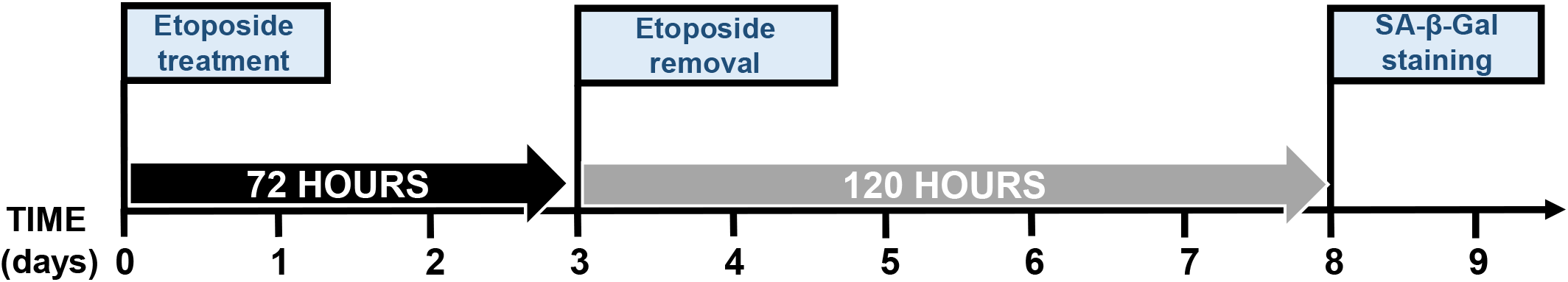
DNA damage-induced growth arrest and senescence of cancer cells induced by etoposide treatment in vitro. Cancer cells were exposed to 5 μM etoposide for 72 hours. The compound was subsequently removed and the cells were cultured with regular growth medium for additional 120 hours. Cells were then fixed in 0.5% glutaraldehyde for 15 min and stained for senescence associated β-Galactosidase as described in Krizhanovsky et al., 2008.

**Figure 5–figure supplement 2.**
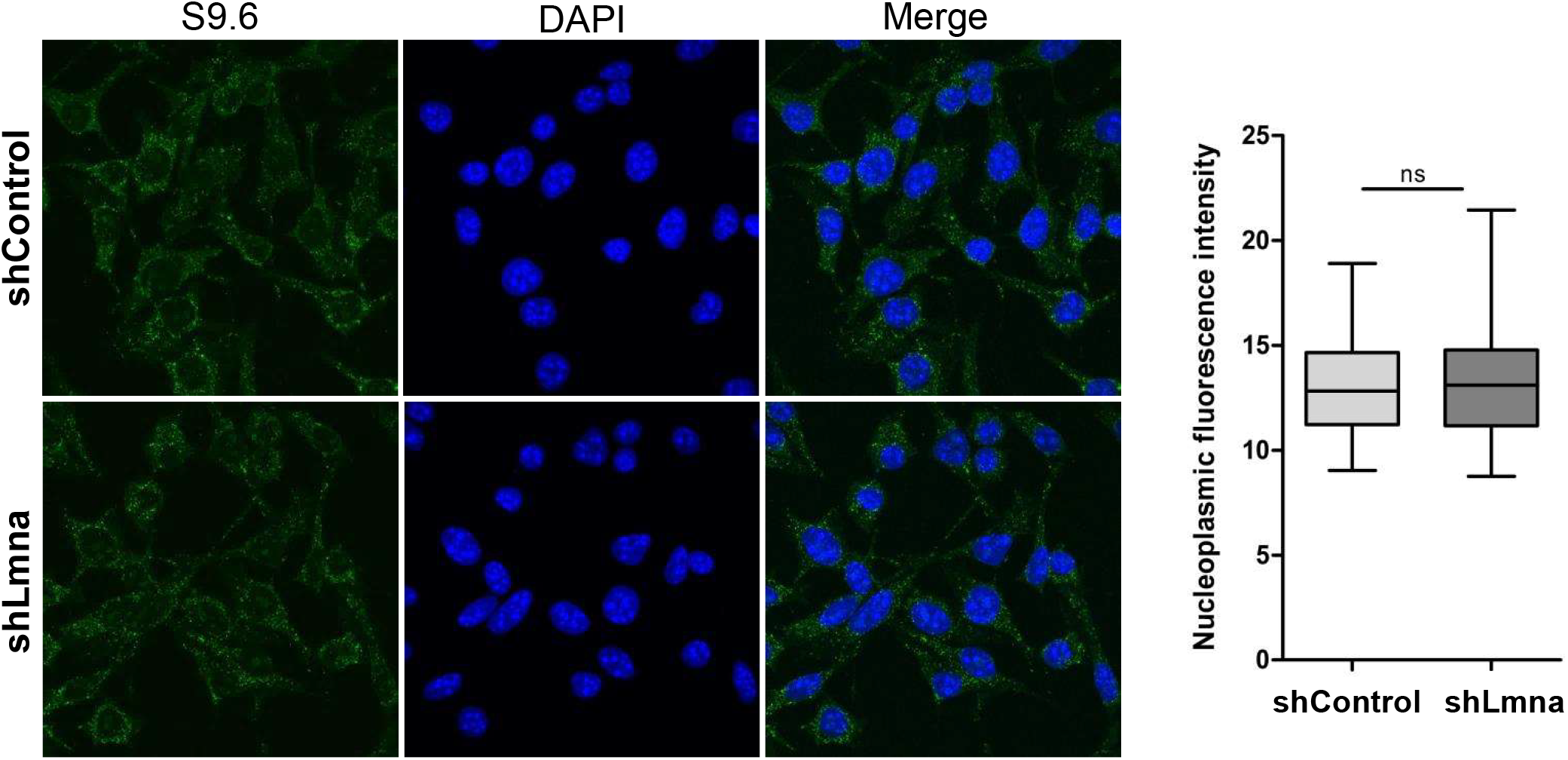
Lamin A/C downregulation doesn’t alter R-loop (S9.6) formation in B16F10 cells. R-loop mapping via S9.6-based anti DNA:RNA hybrid antibody immunofluorescence.

**Figure 5–figure supplement 3.**
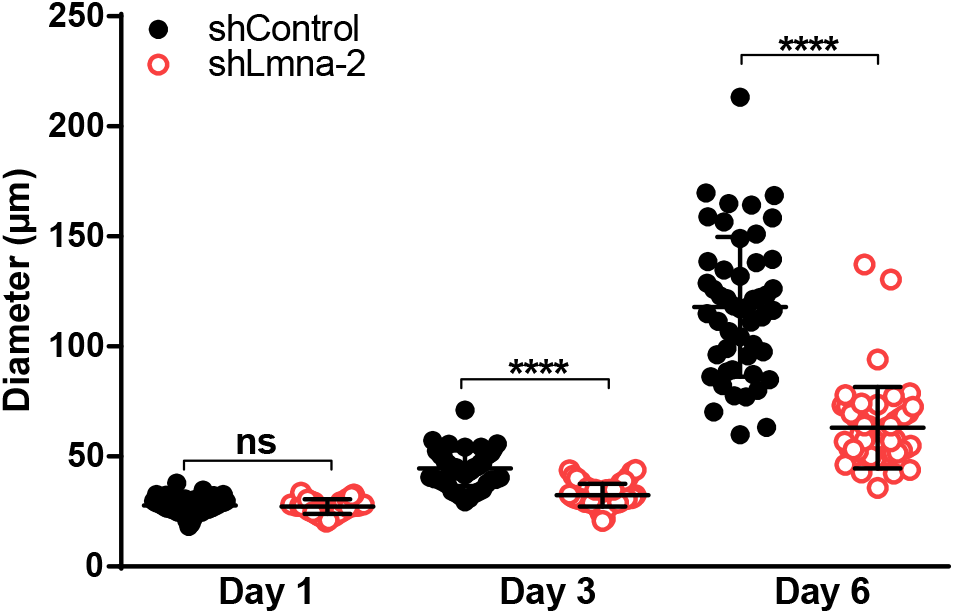
Soft agar colony formation assay. B16F10 (either shControl or shLmna-2 expressing) cells were embedded in soft agar and their growth was monitored for 6 days. Spheroid diameter was measured at days 1, 3, and 6 after seeding. n = 50 cells per group. Values represent the mean ± SEM. The experiment shown is representative of two. ****p < 0.0001.

**Figure 5–figure supplement 4.**
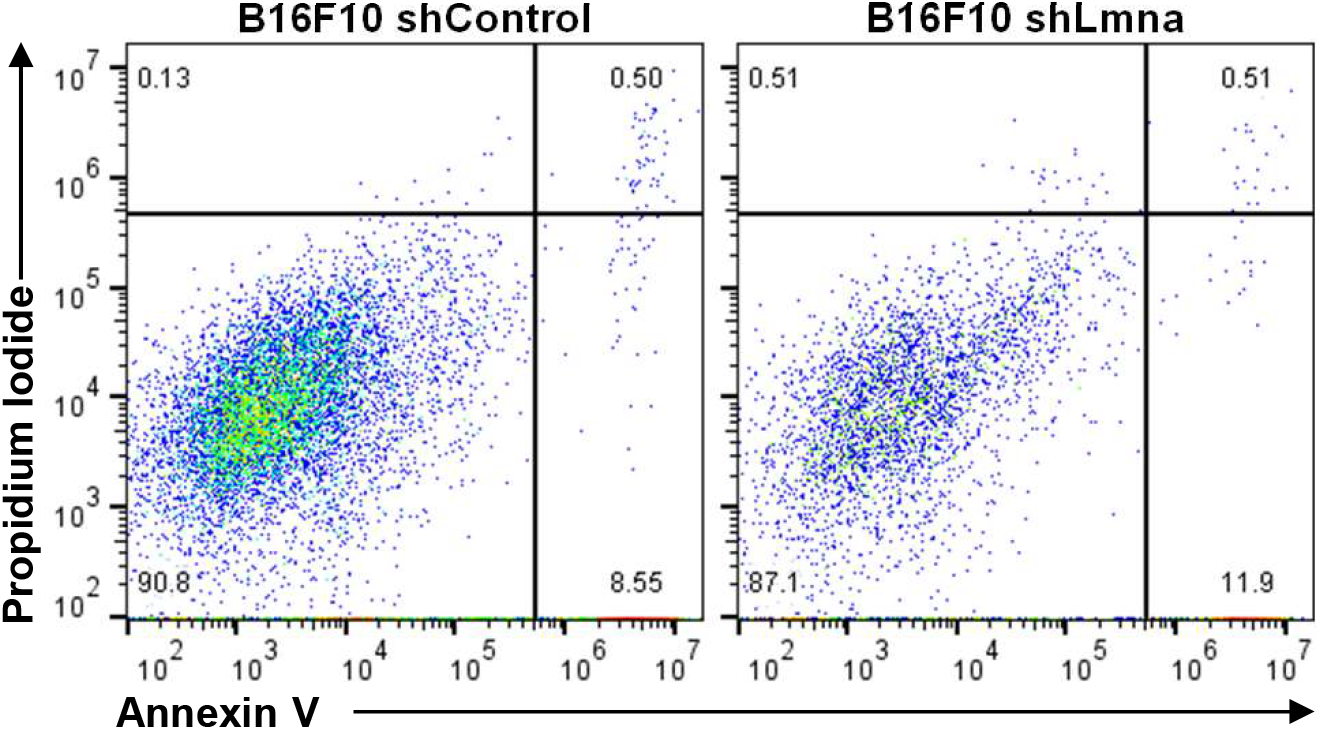
Representative flow cytometry data to illustrate gating and quadrant strategy for measurement of apoptosis. Flow cytometry plots showing annexin V (X-axis) and propidium iodide (Y-axis) staining of B16F10 (either shControl or shLmna expressing) cells derived from 3D colonies grown in soft agar and extracted at day 6. The right lower quadrant represents annexin V positive/propidium iodide (PI) negative staining indicating early apoptosis. The right upper quadrant represents high annexin V and high PI staining indicating late apoptosis and the left upper quadrant represents low annexin V and high PI staining indicating necrosis. The left lower quadrant indicates viable cells.

**Figure 7–figure supplement 1.**
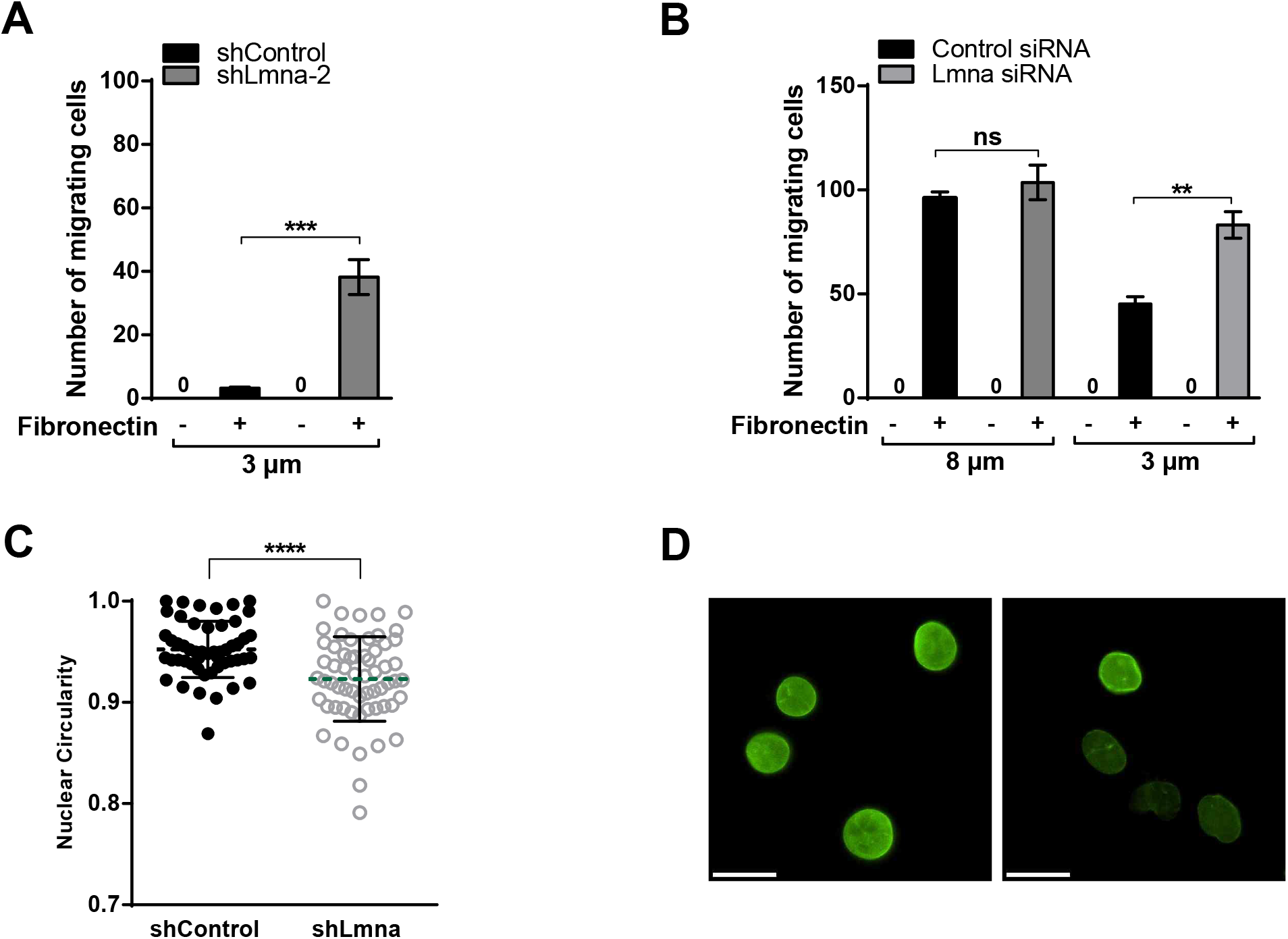
Downregulation of lamin A/C increases E0771 breast carcinoma cells squeezing through small rigid pores and alters nuclear shape. (A) Haptotactic of E0771 shControl vs shLmna-2 through 3 μm pores transwell filters coated (+) or uncoated (−) with Fibronectin (1.5 μg/ml) for 4 h. *** p = 0.0002. (B) Haptotactic migration of E0771 Control and Lmna siRNA cells (72 h post transfection), through 8 or 3 μm pores transwell filters coated (+) or uncoated (−) with fibronectin (1.5 μg/ml) for 4 h. Values represent the mean ± SEM of five fields of view in each experimental group. Results shown are from a representative experiment of three. ***p = 0.0018. (C) Nuclear circularity of E0771 shControl and shLmna cells spread on a bEnd.3-derived basement membrane. ****p <0.0001. (D) Immunostaining of lamin A/C (green) in E0771 shControl and shLmna cells. Scale bar represents 20 μm.

**Figure 7–figure supplement 2.**
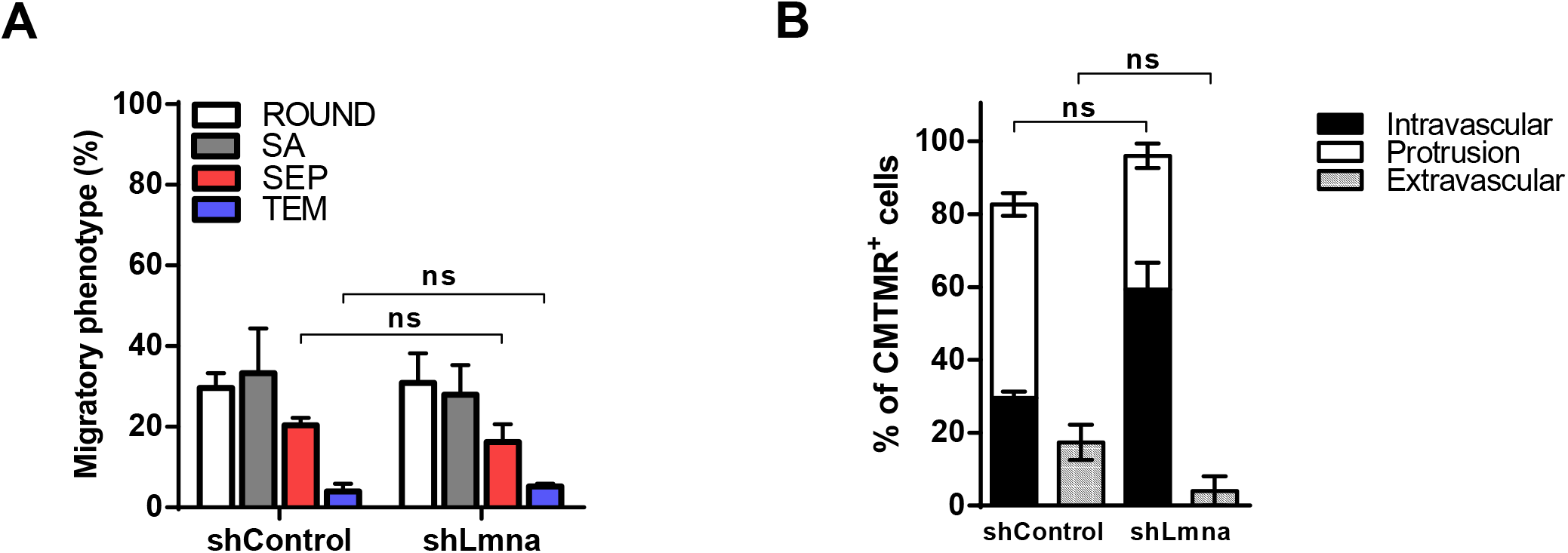
In vitro and in vivo breast carcinoma crossing of endothelial barriers is not enhanced by lamin A/C downregulation. (A) Migratory phenotypes of E0771 breast carcinoma cells TEM. Distinct tumor cell categories (referred to as migratory phenotypes) taken from time lapse videomicroscopy segments of individual E0771 cells: round, spread above (SA), forming sub endothelial pseudopodia (SEP), and completing transendothelial migration (TEM). Values represent the mean ± SEM of three fields in each experimental group. (B) Percentage of E0771 shControl and shLmna cells in a volume of 5×10^9^ μm^3^ of the left lung lobe (3 days after injection). Values are mean ± SEM of 3 different lung’s sections fields of view in each experimental group.

**Figure 7–figure supplement 3.**
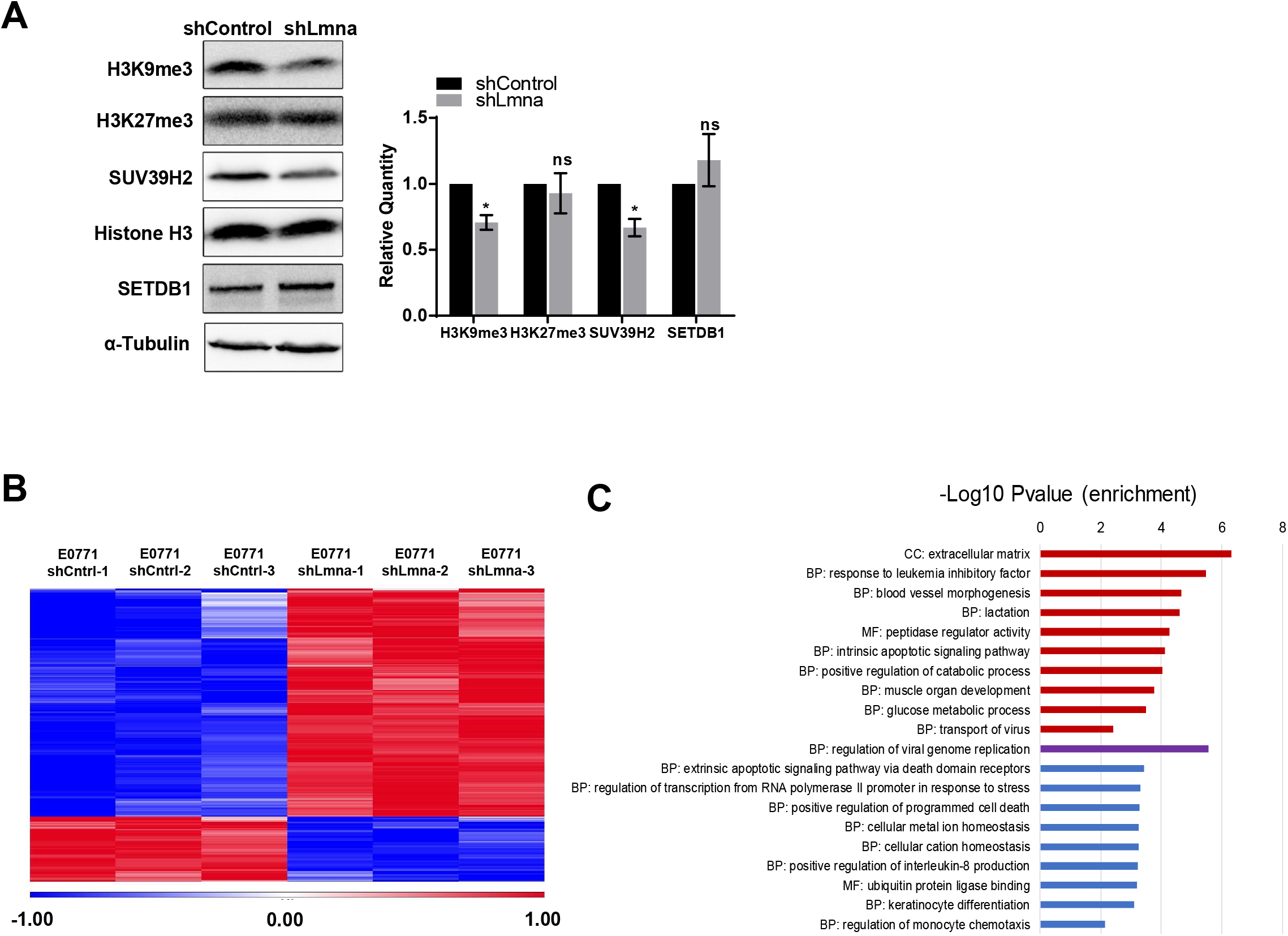
Lamin A/C downregulation reduces heterochromatin content and alters gene transcription. (A) Equal protein amounts from E0771 shControl or shLmna cells, separated by SDS-PAGE and analyzed for the indicated proteins by Western blot analysis. The bar graph represents the mean levels of H3K9me3, H3K27me3 and SUV39H2 normalized to Histone H3 and of SETDB1 normalized to α-Tubulin ± SEM of at least four independent experiments. *p < 0.05. (B) Heatmap based on clustering of 290 differentially expressed genes. Red and blue bars represent positive and negative changes, respectively, and the intensity of the color represents the standardized log2 expression level. (C) Gene ontology (GO) enrichment analysis of the top differentially downregulated (blue) and upregulated (red) genes in E0771 shLmna cells. Biological Process (BP), Molecular Function (MF) and Cellular Component (CC).

**Figure 7–figure supplement 4.**
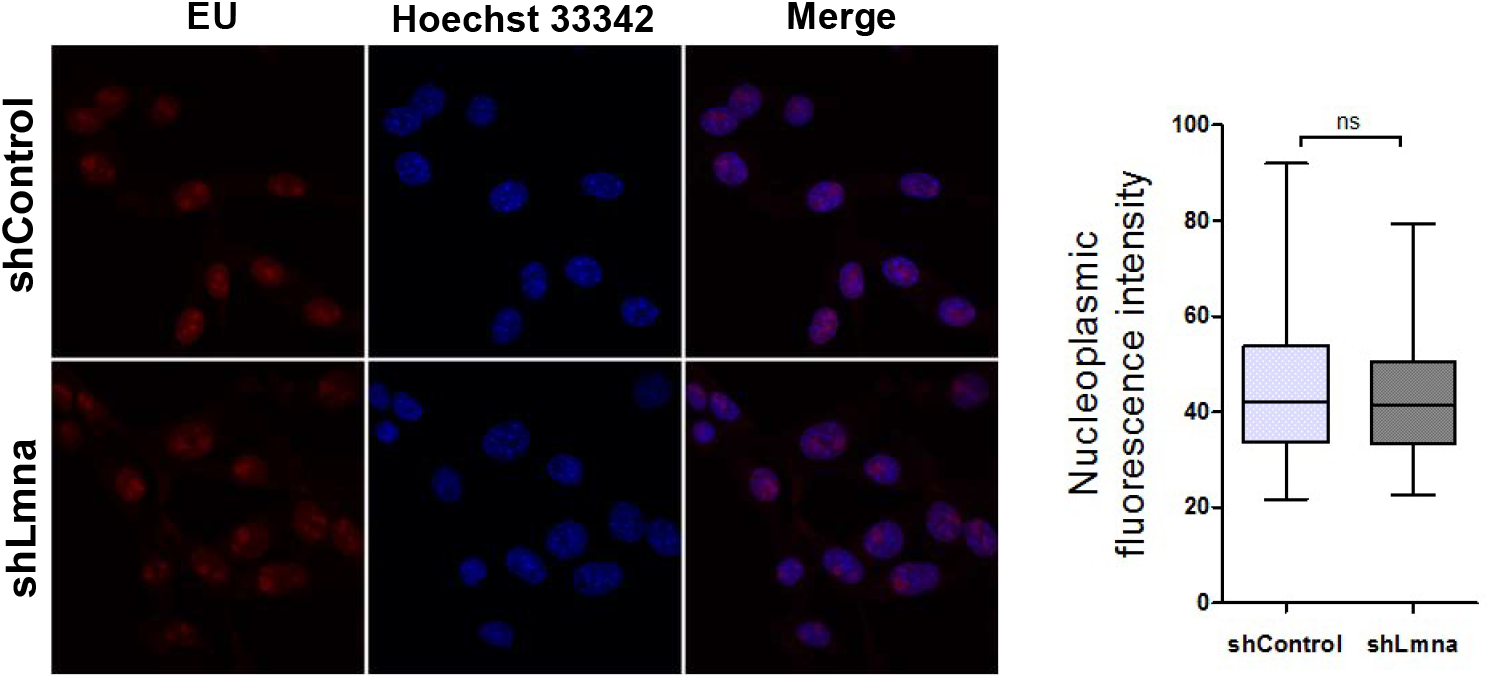
Fluorescence microscopy imaging of 5-ethynyl uridine (EU) incorporation (red) and Hoechst 33342 (blue) in E0771 shControl and shLmna cells. Cells were grown with 1 mM EU for 1 h. The cells were fixed, permeabilized and treated with Alexa Fluor 594 azide. Nucleoplasmic fluorescence intensity of the EU staining was measured using ImageJ. Data from 3 independent experiments are shown in the boxplot.

**Figure 7–figure supplement 5.**
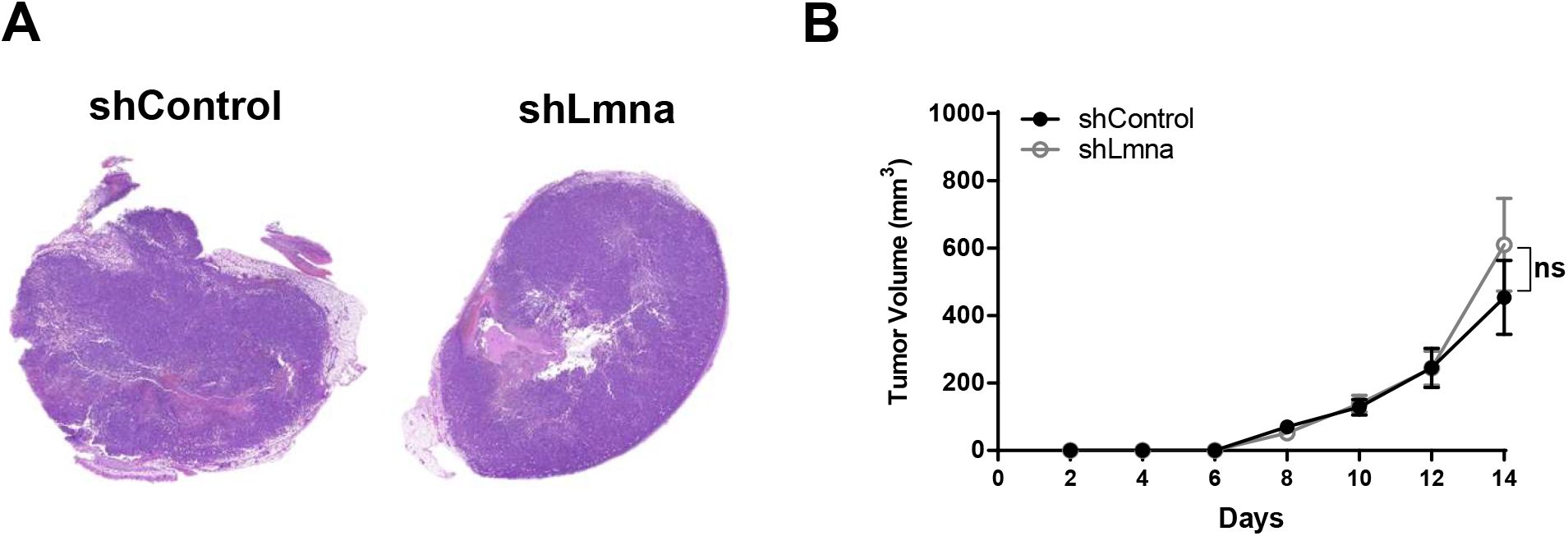
Primary E0771 growth in the mammary fat pad or in the skin. (A) Histology sections of breast primary tumors. A suspension of 10^4^ E0771 (either shControl or shLmna expressing) cells in 50 μl of Matrigel^®^ Matrix mixed with PBS (at 1:1 v/v) was inoculated in the mammary fat pad of recipient mice. 14 days later, animals were euthanized, tumors extracted, and fixed in 4% PFA. (B) Number of E0771 shControl and shLmna present in the lungs of recipient mice 3 hours, 3 and 7 days after i.v. injection. n = 3 for each experimental group. Data are mean ± SEM. The experiment shown is representative of three. (B) 10,000 E0771 shControl and shLmna cells were implanted subcutaneously in flank of C57BL/6 mice. Tumor growth was assessed every other day for 14 days post implantation. The experiment shown is representative of two.

**Figure 7–figure supplement 6.**
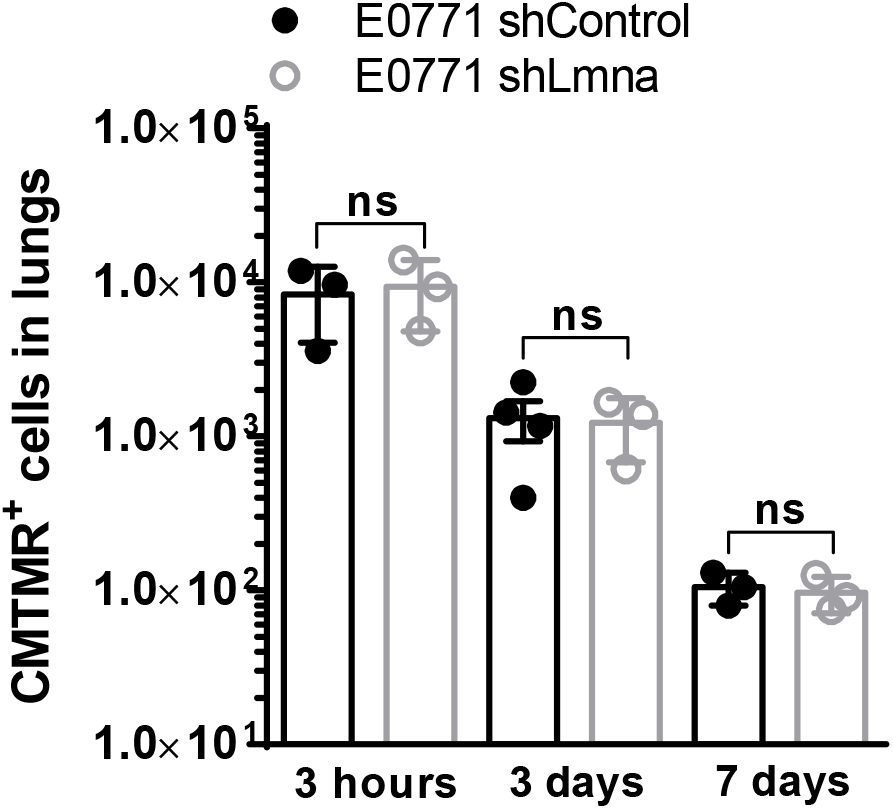
E0771 accumulation in recipient lungs. Number of E0771 shControl and shLmna recovered in the lungs of recipient mice 3 hours, 3 and 7 days after i.v. injection of 20,000 cells. n = 3 or 4 for each experimental group. Data are mean ± SEM. The experiment shown is representative of two.

